# Local Adaptive Mapping of Karyotype Fitness Landscapes

**DOI:** 10.1101/2023.07.14.549079

**Authors:** Richard J Beck, Noemi Andor

## Abstract

Despite its critical role in tumor evolution, a detailed quantitative understanding of the evolutionary dynamics of aneuploidy remains elusive. Here we introduce ALFA-K (Adaptive Local Fitness landscapes for Aneuploid Karyotypes), a novel method that infers chromosome-level karyotype fitness landscapes from longitudinal single-cell data. ALFA-K estimates fitness of thousands of karyotypes closely related to observed populations, enabling robust prediction of emergent karyotypes not yet experimentally detected. We validated ALFA-K’s performance using synthetic data from an agent-based model and empirical data from in vitro and in vivo passaged cell lines. Analysis of fitted landscapes suggests several key insights: (1) Whole genome doubling facilitates aneuploidy evolution by narrowing the spectrum of deleterious copy number alterations (CNAs); (2) Environmental context and cisplatin treatment significantly modulate the fitness impact of these alterations; (3) Fitness consequences of CNAs are contingent upon parental karyotype; and (4) Chromosome mis-segregation rates strongly influence the predominant karyotypes in evolving populations.

## Introduction

Losses and gains of entire chromosomes or large sections thereof, known as aneuploidy, are a defining feature of solid tumors[1, 2]. This aberrant state arises from an ongoing dynamic process termed chromosomal instability (CIN), which stems from diverse mechanisms including mitotic segregation errors, replication stress, and structural chromosome damage[3]. While CIN can also manifest as structural aberrations, this work focuses primarily on whole-chromosome CIN and the resulting numerical aneuploidy state[3], as these whole-chromosome changes affect more of the cancer genome than any other genetic alteration[4]. Present in an estimated 90% of solid tumors, aneuploidy modifies the copy number of many genes, thereby altering cellular phenotype through correlated changes in RNA expression and protein production[5, 6]. Aneuploidy provides a substrate for tumor evolution[1, 7, 8], often enriching chromosomes with oncogenes while deleting those with tumor suppressor genes[9].

The factors which explain aneuploidy patterns in cancers are not limited to the density of driver or suppressor genes on a particular chromosome. Aneuploidy is usually detrimental to cell fitness [10], in the first instance due to proteins such as P53 which cause apoptosis or cell cycle arrest in response to chromosome missegregations [11]. According to the gene dosage hypothesis, aneuploidy can also reduce fitness by upsetting the balance of protein levels within cells: leading to negative effects such as impaired formation of stoichiometry dependent protein complexes, or protein aggregates that overwhelm protein quality-control mechanisms [10, 11]. In addition to these intracellular effects, the environmental context plays a role in sculpting karyotype [4], since the specific pressures of an environment will determine whether the fitness advantages of a particular copy number alteration (CNA) outweigh the costs. Evidence for the role of environment in determining karyotype includes the selective advantage of particular karyotypes under stressful conditions in yeast [5] and distinct patterns of aneuploidies between cancer types [12, 13, 14]. Genomic context also plays a role in sculpting karyotype because a given CNA may only be favorable if other mutations or CNAs are already present within the cell. Support for the role of genomic context comes from findings that CNAs lacking independent prognostic value can predict survival in combination [15], as well as from the reproducible temporal ordering of CNAs in patient-derived xenograft and organoid models [16, 17].

Aneuploidy remains difficult to study, for reasons which include the difficulty of experimentally inducing aneuploidy and the difficulty of distinguishing the effects of aneuploidy from those of chromosomal instability, the process which causes aneuploidy [4]. *In silico* models will be an important tool to further our knowledge of aneuploidy. Gusev and colleagues developed the first model describing whole-chromosome missegregations [18, 19]. This model laid the mathematical foundations for describing segregation errors and explained patterns of aneuploidy in experimental data as a consequence of variable chromosome missegregation rate. A limitation of this model was that a fitness landscape defining the effect of aneuploidy on cell fitness was not considered, beyond a constraint that cells losing all copies of any chromosome were not considered viable. This limitation was later addressed by others who assumed that the fitness effect of changing the copy number of a particular chromosome was dependent on the number of oncogenes or tumor suppressor genes expressed on that chromosome [20, 21]. In these models cell fitness could be increased by gaining additional copies of chromosomes with many oncogenes, or losing copies of chromosomes with many tumor suppressor genes. These models predicted an optimal missegregation rate (in the sense of minimising total cell death) that matched experimental observations, and resulted in a near-triploid karyotype that is frequently observed in tumour cells. More recent work explored alternative fitness assumptions, considering not only driver gene density but also stabilizing selection based on overall gene abundance or a hybrid of both approaches, while using simulations to infer CIN rates [22]. All of these previous models of aneuploidy fail to account for how environmental influences and genomic background impact the relationship between karyotype and fitness. This is perhaps unsurprising, since the vast number of possible karyotypes is challenging enough to map even without the additional variability introduced by these contexts. However, the burgeoning quantity of single-cell copy number data [23, 24] now permits tracking of subclonal evolution at unprecedented resolution, providing the potential to refine our understanding of the relationships between karyotype and cellular fitness.

Fitness landscapes represent a mapping from genome to cellular fitness. Charting these landscapes for karyotypes is particularly challenging due to the vast number of possible states - over 10^19^ karyotypes when considering up to eight copies per chromosome. Here, we introduce ALFA-K, the first method to directly chart local regions of karyotype fitness landscapes from single-cell copy number data. After validating ALFA-K with synthetic data from an agent-based model of chromosome missegregations, we further confirmed its predictive accuracy using empirical data from P53-deficient cell lines showing extensive subclonal evolution. Our analysis yielded several novel insights, highlighting roles for whole genome doubling, *in vivo* selection pressures, cisplatin treatment, and chromosome missegregation rates in shaping the topology of karyotype fitness landscapes and cellular evolutionary trajectories.

## Results

### ALFA-K predicts karyotype evolution of variable speed and complexity

We developed a framework to infer **A**daptive **L**ocal **F**itness landscapes for **A**neuploid **K**arotypes (ALFA-K). The method utilizes longitudinal data from evolving cell populations (Fig. 1A), where single cells are analyzed via sequencing to determine their specific karyotypes. This allows for tracking temporal changes in the frequency of common karyotypes within the population (Fig. 1B). ALFA-K assumes that fitness is intrinsically determined by karyotype, equating to the net growth rate, and that this fitness landscape is static over the observation period (conceptualized in Fig. 1C). Based on this, fitness estimates for common karyotypes are derived directly from their observed frequency changes over time. These initial estimates are then extended to rarer, related karyotypes through Gaussian process regression, thereby constructing an inferred local fitness landscape covering the explored karyotypic space (Fig. 1D). The framework uniquely accounts for the fact that the fitness consequences of missegregation (MS) can vary depending on the specific karyotype background of the missegregating cell (Supplementary Section 1).

**Figure 1.**
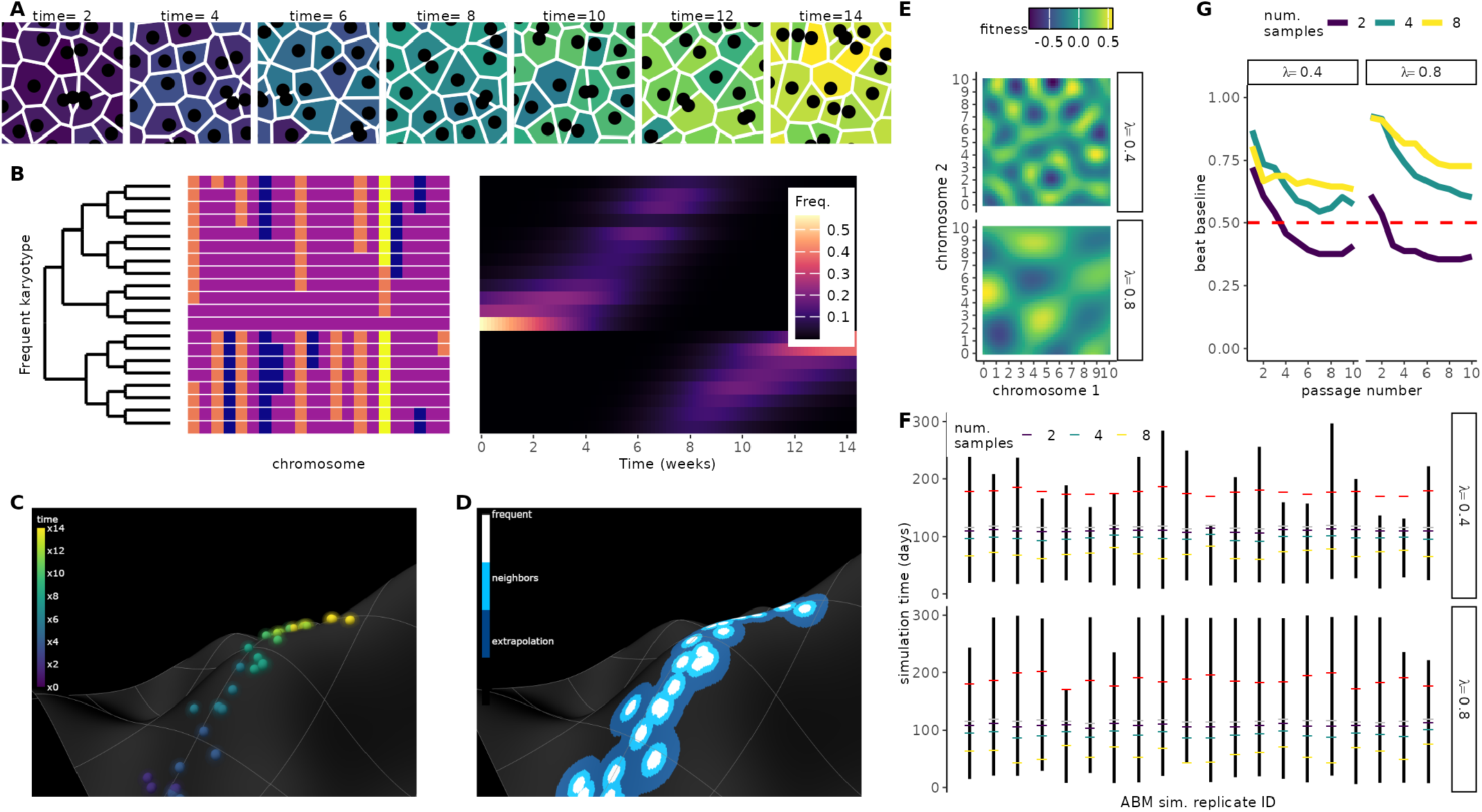
Inferring aneuploid fitness landscapes with ALFA-K. Conceptual overview. (**A**) Schematic representation of an evolving cell population passaged longitudinally over multiple timepoints (panels). Individual cells are colored based on the timepoint at which their specific karyotype first emerged in the population. (**B**) Karyotypes (rows) are determined from single-cell sequencing data (Supplementary Section 4.1). The left heatmap displays inferred chromosome copy numbers (fill color) for various detected karyotypes. In the right heatmap, the frequency (fill color) of distinct karyotypes (y-axis) across different timepoints (x-axis) is shown. (**C**) Conceptual visualization of karyotype evolution on a fitness landscape. Each point represents a unique karyotype, positioned according to a 2D projection of its high-dimensional state (x and y axes), with fitness indicated by height (z-axis). Points are colored by their time of first emergence, corresponding to panel A. (**D**) Using the same representation as C, but highlighting instead the region where fitness estimates are made by the pipeline. Fill color within the charted region indicates the stage of the inference process used to estimate fitness for that specific karyotype (e.g., direct frequency-based estimation vs. Gaussian process regression, see Supplementary Section 1). **Validation of forecasting performance**. (**E**) Two example Gaussian-random-field (GRF) fitness landscapes illustrate how increasing the wavelength (*λ*) alters topology. (**F**) Overview of ABM sampling strategy. Agent-based simulations incorporating MS-driven karyotype changes were run on the GRF landscapes in (E); the resulting karyotype counts served to train and validate ALFA-K. Black bars indicate the longest continuous fitness-increasing interval per simulation; colored ticks mark the time windows used for training; grey dots indicate the final training timepoint, and red dots mark the prediction horizon. (**G**) Fraction of ABM simulations whose forecasts outperform a Euclidean “no-evolution” baseline (see Supplementary Section 3.1), evaluated on unseen training data after excluding landscapes with negative CV scores.

We first validated ALFA-K on synthetic datasets. We set up agent-based model (ABM) simulations evolving on Gaussian random-field fitness landscapes (Fig. 1E), then trained and evaluated ALFA-K performance on simulated data across a period of active population evolution (Fig. 1F). Accuracy increased on smoother terrains and when more longitudinal passages were available, whereas systematic error stemmed mainly from inaccurate fitness estimates for well-sampled (“frequent”) karyotypes (Fig. S2). ABM simulations using fitted fitness landscapes as inputs reproduced clonal dynamics (Figs. 1G, S4), supporting subsequent application to longitudinal single-cell experiments.

To evaluate ALFA-K on experimental data, we fitted the model to longitudinal single-cell karyotype profiles from two breast-cancer cell lines and four patient-derived xenograft (PDX) transplant series cultured with or without cisplatin [23] (Fig. 2A, see Supplementary Section 4.1 for details). Because ground-truth fitness landscapes are unknown for experimental data, model accuracy was quantified with CV scores. Here, a <u>trajectory</u> is the ordered sequence of passages obtained from a single lineage (e.g. *A*→ *B* →*C*) and its nested sub-sequences (e.g. *A* → *B, B* → *C*); each trajectory yields one ALFA-K landscape. Across the six independent lineages this definition produced **275** candidate trajectories. In line with *in silico* tests, CV scores generally increased with the number of training passages used (Fig. 2B). ALFA-K landscapes obtained on trajectories of size four or more resulted in a median CV score of 0.48 (cluster-bootstrap 95% CI 0.11–0.57). Across all fitted landscapes with a positive CV score, ALFA-K returned fitness estimates for 272,317 unique karyotypes.

**Figure 2.**
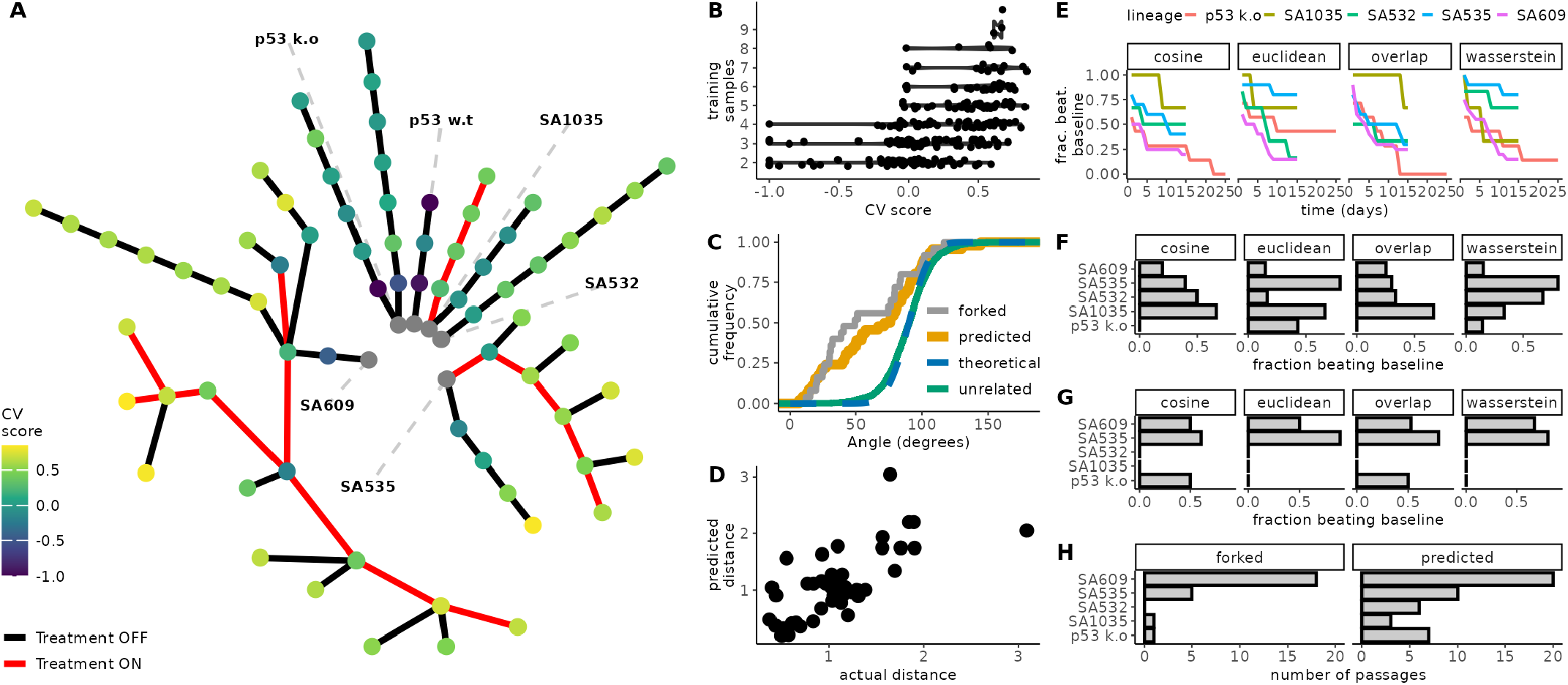
Validation of ALFA-K on longitudinal single-cell data. (**A**) Lineage graph showing experimental passages (nodes) coloured by ALFA-K CV score, used here as a proxy for model fit in the absence of ground truth fitness. Red edges indicate intervals of cisplatin treatment. To select inputs for forecast validation (C-G), we defined sublineages as any passage plus a preceding history; for each potential endpoint, the history length and frequent clone threshold maximizing the CV score were chosen. Sublineages with CV score ≤ 0 were excluded, yielding 35 sublineages for analysis. (**B**) CV score generally improves with the number of passages used for training. (**C**) Cumulative frequency distribution functions (CDFs) of angle metrics comparing forecasts to ground truth. ALFA-K forecasts (predicted) show similar directional accuracy to comparisons between sister passages (forked). Both are significantly more aligned with the true direction than the random-orientation null (theoretical), which is matched by comparisons between unrelated lineages. (**D**) Predicted vs. observed Wasserstein distances between consecutive passages. (**E**) Fraction of the 35 selected sub-lineages where ALFA-K forecasts outperform a static “no-evolution” baseline, plotted against time since training for all metrics. (**F**) Fraction of ALFA-K forecasts outperforming the static baseline at the specific forecast horizon corresponding to the next measured passage. (**G**) Fraction of sister passages which “beat baseline”, evaluated on available forked passages. (**H**) Number of passages available for metric evaluation corresponding to F-G.

We then tested ALFA-K’s ability to forecast the karyotype distribution of the subsequent passage. We retained fits with positive cross-validation score and at least one downstream passage, keeping only the highest-scoring fit per final timepoint so that each defined a distinct test instance (N=35 fitted lineages). We used the angle metric (a measure of similarity between two evolutionary trajectories, where an angle of 0 degrees indicates identical paths of change) to evaluate predictive performance, calculating the angle between observed evolutionary changes and either ALFA-K forecasts or experimental sister passages. Both sister-passages (median-of-lineage-medians 79.0^*°*^, cluster bootstrap *p* = 0.0076) and ALFA-K predictions (median-of-lineage-medians 71.2^*°*^, cluster bootstrap *p <* 0.0001), were significantly more aligned than random orientations. Comparisons between unrelated lineages followed the null distribution (median-of-pair-medians 90.4^*°*^, parametric double-cluster bootstrap *p* = 0.80), confirming that the model captures lineage-specific evolutionary signal (Fig 2C). Furthermore, ALFA-K recovered the overall scale of evolutionary change; predicted Wasserstein distances between consecutive passages correlated well with the observed distances (Spearman *ρ* = 0.68, Pearson *r* = 0.72, Fig 2D).

Next, we compared ALFA-K forecasts to a static “no-evolution” baseline, where the predicted state is simply the last observed state. The fraction of sub-lineages where ALFA-K outperformed this baseline was highest immediately after training and declined over 15–25 days (Fig 2E). Evaluating performance specifically at the forecast horizon corresponding to the next measured passage, we compared ALFA-K’s baseline-beating rate to that achieved by using experimental sister passages as predictors (Fig 2F, G). Although sister-passage predictions outperformed the static baseline in 56% of cases, more frequently than ALFA-K forecasts (in 36%), this comparison requires caution as the lineage sets differed; ALFA-K evaluation was restricted to sub-lineages with high CV scores, while the sister-passage comparison depended on the availability of experimentally forked passages (Fig 2H). Overall, the rates at which both methods surpassed the baseline were modest, underscoring the difficulty in forecasting subtle clonal shifts over these short intervals.

These results suggest that ALFA-K captures directional and quantitative trends in karyotype evolution, offering predictive power that approaches replicate-passage comparisons, particularly in the near term. However, the modest rates at which forecasts outperformed the static baseline (Fig 2E-G) highlight the inherent challenge of making accurate, fine-grained predictions of clonal population dynamics over typical experimental timescales.

### ALFA-K Predicts the Emergence of Novel Karyotypes

Prior work by Salehi et al. [23] demonstrated that the dynamics of karyotype-defined subpopulations can be extrapolated over time, but their method was limited to karyotypes already observed. In contrast, we asked whether ALFA-K could forecast the emergence of previously unobserved karyotypes—i.e., genotypes that had not yet appeared but could arise based on their fitness and location in the karyotype space.

To evaluate the predictive performance of ALFA-K for previously unobserved karyotypes, we focused on the same subset of 35 fitted karyotype landscapes used in earlier predictive analyses (Fig. 2C–F), selected for their availability of future time points that allow assessment of novel karyotype emergence. Clonal interference frequently arises in these landscapes, where multiple fit karyotypes co-exist and compete for dominance over time. As shown in Figure 3A, these dynamics can obscure which karyotypes will ultimately expand, especially when several are predicted to have similarly high fitness. Capturing the effects of such competition is therefore central to ALFA-K’s ability to correctly forecast emergent karyotypes. The selected lineages were distributed across five cell lines as follows: p53 k.o – 7; SA1035 – 3; SA532 – 6; SA535 – 6; SA609 – 13.

**Figure 3.**
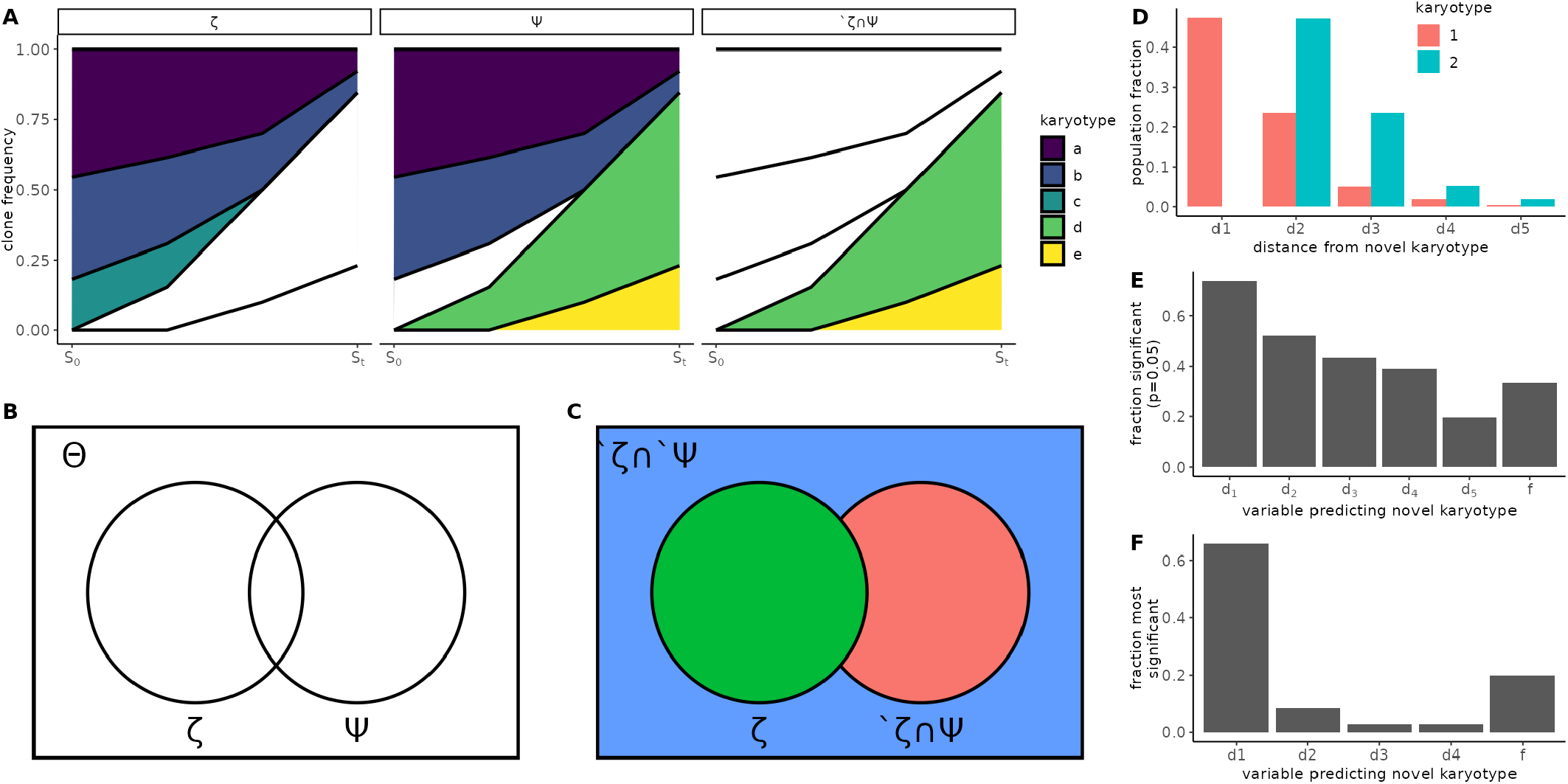
ALFA-K predicts emergence of novel karyotypes. (**A**) Clonal interference—simultaneous expansion of multiple advantageous karyotypes that compete and slow one another’s fixation—is illustrated for 5 example karyotypes. Clone group (columns) is categorized as illustrated in B-C. *S*_0_ and *S*_*t*_ indicate present and future sample timepoints, respectively. (**B**) Venn diagram representing all karyotypes in the fitted landscape (Θ), the subset observed in the latest sample (*ζ*), and the subset that will be present in a future sample (Ψ). (**C**) Θ is separated into 3 disjoint subsets. The aim is to predict ^*′*^*ζ* ∩ Ψ. (**D**) Distance feature vector assigned for two example karyotypes: **k**_1_, **k**_2_ ∈*′ ζ* ∩ Ψ. *d*_*i*_ is the fraction of karyotypes in *ζ* that are *i* missegregations away from **k**_1_/**k**_2_. E-F) Comparing the contribution of *f* (the ALFA-K estimated karyotype fitness) to that of other variables as predictors of novel karyotypes **k**^∗^ ∈ ^*′*^ *ζ* ∩Ψ. Fitness estimates used here are computed using data prior to the point of emergence being predicted to avoid look-ahead bias. (**E**) Fraction of fitted lineages in which each variable was significant (*P <* 0.05). (**F**) Fraction of fitted lineages in which each variable was most significant. Significance (E-F) was determined by the Wald test P-value of the test statistic for each fitted parameter.

Let Θ denote the set of karyotypes with fitness estimates from ALFA-K, *ζ* the subset of Θ observed in a given longitudinal sample. Then we would like to predict Ψ (the subset of Θ that will be present in a future sample) (Fig.3A-B). In particular we wish to predict which new karyotypes will emerge in the next sample, ^*′*^*ζ* ∩Ψ (Fig.3C). The probability of any novel karyotype actually emerging presumably depends both on its fitness and its number of neighbours in the preceding generation. Therefore for each member of ^*′*^*ζ* we computed the fraction of karyotypes in *ζ* that were between 1-5 missegregations distant (Fig.3D). These were used as variables which, together with the fitness estimate (*f*), were used to predict whether the karyotype would emerge. For prediction we used binomial logistic regression, then assessed whether each predictor variable was significantly correlated with the response variable. As expected, the fraction of *ζ* that were distance-1 neighbours (*d*_1_) was a very significant predictor of novel karyotype emergence (Fig.3E-F). The fitness estimates from ALFA-K also contributed significantly to the prediction, being significant in 15/45 tests (Fig.3E) and the most significant predictor in 7/45 tests (Fig.3F). These results indicate that ALFA-K fitness estimates can help predict emergence of new karyotypes.

### Karyotypic background determines fitness effects of CNAs

We set out to quantify how experimental context (*in vitro* vs. PDX) and cisplatin exposure reshape the fitness landscapes inferred with ALFA-K. To ensure statistical independence, we implemented a bootstrap procedure, repeatedly sampling sets of evolutionary trajectories with no shared passages (Figure 4A). For each of 200 bootstrap replicates we re-estimated all model coefficients, and deemed an effect significant (*p <* 0.05) when its 95% bootstrap confidence interval did not include zero. For each frequent karyotype in every fitted landscape we calculated the 44 one-MS-step fitness effects (Δ*f*) (Fig. 4B). Visualizing the raw Δ*f* distributions for a typical bootstrap replicate revealed only minor differences in the median but substantial broadening of the distribution tails (Fig. 4C). To quantify the visual pattern, we used two GLMMs—one Gaussian on the log-variance of each karyotype’s Δ*f* profile, the other log-Gamma on the absolute Δ*f* across all karyotypes and landscapes. These approaches examine complementary aspects of the same underlying landscapes: the log-variance captures how widely fitness effects vary within lo-

**Figure 4.**
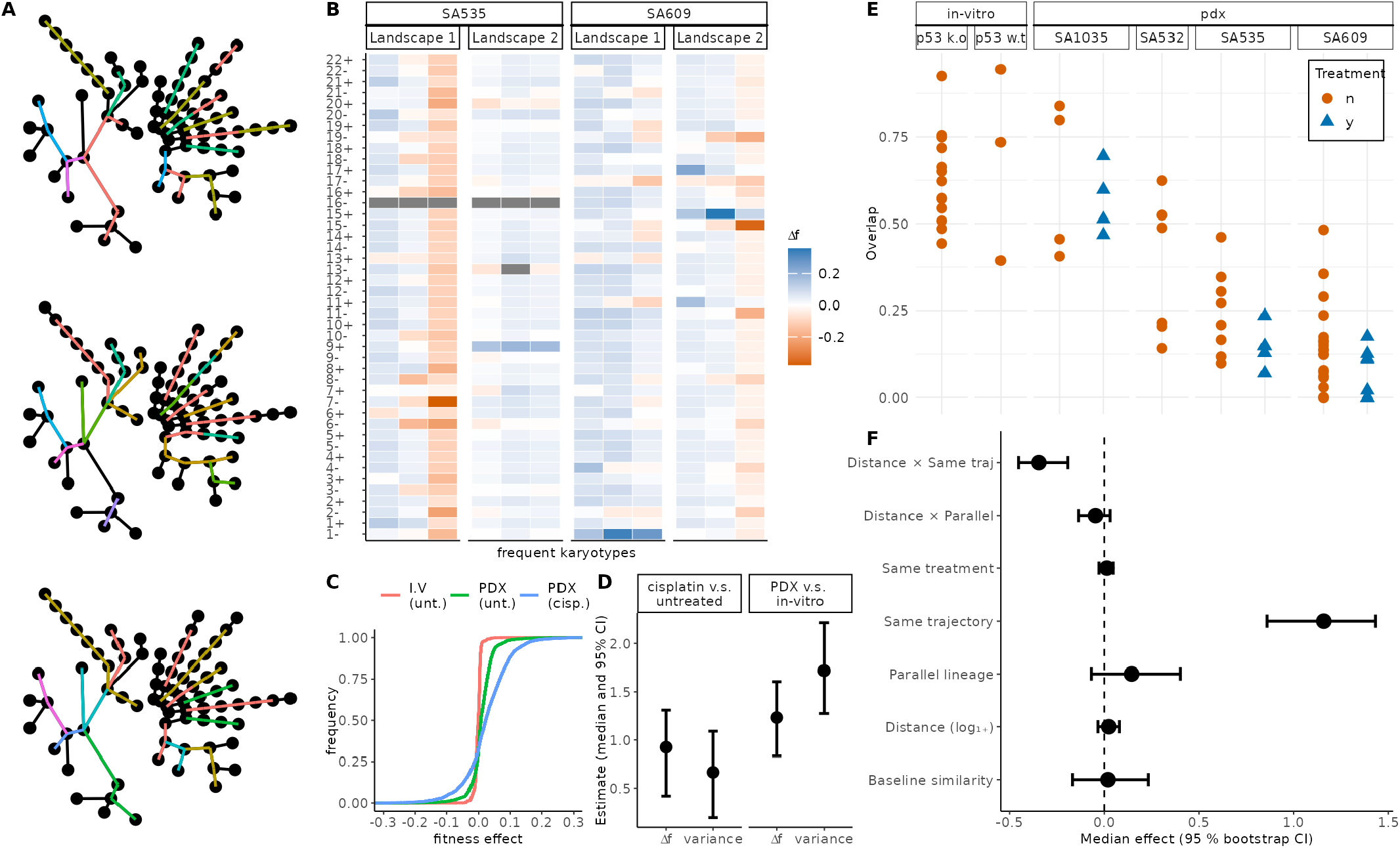
Parent cell karyotype and treatment context shape CNA fitness effects. (**A**) Radial dendrograms of all sequenced passages. Coloured segments indicate different sets of non-overlapping evolutionary trajectories used within different bootstrap replicates. (**B**) For each karyotype, the fitness change associated with gaining or losing each chromosome was predicted by ALFA-K. The resulting vectors (Δ*f* profiles) for sample karyotypes are shown in each column. (**C**) Empirical cumulative-distribution functions of Δ*f* for the three analysed conditions, from a single bootstrap replicate. (**D**) Estimated effects of experimental context and treatment on the fitness landscape. Points show the median coefficient estimates from 200 bootstrap replicates, with bars indicating the 95% confidence intervals. Effects are shown for both the magnitude of fitness changes (Δ*f*) and their variance. (**E**) Abundance-overlap coefficients between consecutive passages, plotted by cell line, colored and shaped by treatment. Lower overlap signifies larger compositional changes from one passage to the next. (**F**) Factors influencing the Pearson correlation of Δ*f* profiles between karyotype pairs. The plot shows the median coefficient estimates (points) and 95% confidence intervals (bars) from a bootstrapped linear mixed model. The “baseline” estimate represents the similarity for karoytypically identical, cisplatin-treated karyotype pairs from different cell lines. The slope term (Distance) quantifies how correlation changes with (log transformed) karyotypic distance, while its interaction coefficients indicate how that distance-decay is modified by pair type or treatment. All remaining categorical coefficients represent additive shifts from the baseline. Confidence intervals that do not intersect zero mark statistically significant effects (*p* = 0.05).

cal karyotypic backgrounds, while the log-absolute |Δ*f*| reflects the typical size of individual mutational steps. The analysis revealed that the coefficients for both PDX context and cisplatin treatment were positive and significant across all models. The 95% confidence intervals, derived from the distribution of estimates across all bootstrap replicates, were entirely above zero, confirming that both conditions significantly increase fitness effect magnitude and variance (Fig. 4D). Together, the elevated variance and larger individual fitness effects suggest that PDX growth and cisplatin exposure increase the availability of both advantageous and deleterious CNA steps. Stronger fitness effects are expected to accelerate clonal expansions and extinctions, leading to greater shifts in population composition between passages. Consistent with this, clone–composition changes between successive passages were more pronounced under these conditions(Fig 4E): overlap coefficients decline from *in vitro* controls to PDX controls and drop further after cisplatin, indicating that compositional shifts are larger when |Δ*f*| and variance are high.

To quantify how similarity between Δ*f* profiles depends on karyotype distance and experimental context, we fitted a linear mixed model with the Pearson correlation between Δ*f* profiles for pairs of karyotypes as the response variable. Relative to a reference of cisplatin-treated pairs with identical karyotype, drawn from different cell lines—whose correlation was indistinguishable from zero (Fig-4E)—three patterns emerged. First, membership in the same inferred fitness landscape markedly increased correlation. Second, within-landscape correlation decreased sharply with growing karyotype distance, confirming a distance-decay relationship. Third, karyotype pairs that evolved in parallel from the same cell line showed a modest uptick in correlation, but this effect did not reach significance in the bootstrap analysis. No other covariates, including shared treatment background, had detectable influence. Taken together, these results underscore that physical proximity in karyotype space is an important determinant of CNA-fitness profile similarity.

We next asked how whole-genome doubling (WGD) alters both the pace of aneuploidy accumulation and its predicted fitness consequences in our two p53 k.o. lineages. Modeling the passaging trajectories revealed that WGD^+^ populations accumulated more chromosome alterations resulting in a significantly higher level of aneuploidy compared to WGD^−^ populations (Fig. 5A, non-linear saturation model, WGD effect on asymptote Wald test *P <* 10^−6^)). ALFA-K fitness estimates generally increased with the number of chromosome alterations in both WGD^−^ and WGD^+^ populations (Fig. 5B). However, the relationship displayed distinct visual patterns: mean WGD^−^ fitness appeared to rise steeply for the first 2-3 alterations before leveling off, while mean WGD^+^ fitness continued to increase across the broader range of aneuploidy observed in these subclones. Because ALFA-K restricts fitness predictions to karyotypes within two mis-segregations of the observed frequent clones (Methods), the upper bound on “chromosomes altered” (Fig. 5B) reflects how far each sub-population actually explored karyotype space. Finally, the empirical cumulative distributions of one-MS-step fitness effects (Fig. 5C) are shifted toward less deleterious outcomes in WGD^+^ subclones (permutation KS test on fit-level averages, empirical *p* = 0.03), suggesting that whole-genome doubling not only accelerates karyotypic diversification but also reduces the pool of fitness-decreasing aneuploid states.

**Figure 5.**
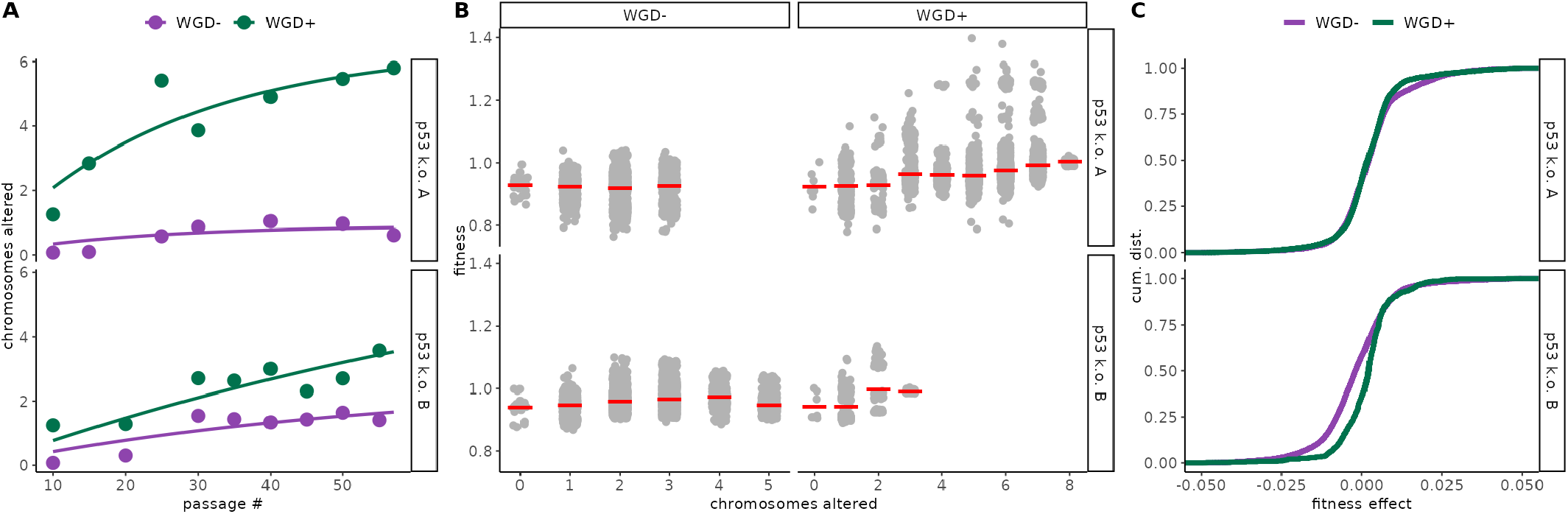
Whole-genome doubling accelerates and reshapes aneuploidy evolution. **A)** Passaging trajectories for p53 k.o. A (top) and p53 k.o. B (bottom): number of chromosomes differing from the euploid state plotted by passage number, colored by WGD status (WGD^−^ = purple, WGD^+^ = green). Curves show fits from a non-linear saturation model accounting for additive effects of WGD status and trajectory on accumulation dynamics (see Supplementary Section 4.2). **B)** ALFA-K–estimated relative fitness versus number of chromosomes altered: light-gray points are individual karyotype estimates; red ticks denote mean within each missegregation bin. **C)** Empirical cumulative distributions of one-MS-step fitness effects for frequently observed karyotypes in each lineage, comparing WGD^−^ (purple) and WGD^+^ (green) subclones.

### Missegregation rate influences karyotype dominance

In most circumstances, the fittest karyotype is expected to eventually dominate the population. However, we asked whether this outcome could be altered simply by changing the missegregation rate. Cells with similar karyotypes that occupy the same fitness peak can be conceptualized as a “quasispecies” [25], and under the quasispecies framework, sufficiently rapid mutation can cause populations to drift off narrow fitness peaks—a phenomenon known as the error threshold [25]. We therefore considered two ways that elevated missegregation might erode dominance (Fig. 6A): first, quasispecies containing many high-fitness karyotypes incur smaller fitness penalties from continual mis-segregation events, because the resulting progeny remain relatively fit; second, low-ploidy quasispecies experience fewer errors per division, allowing for greater population stability. If the fittest karyotypes have few fit neighbors or are of high ploidy, they may ultimately be displaced under high missegregation rates.

**Figure 6.**
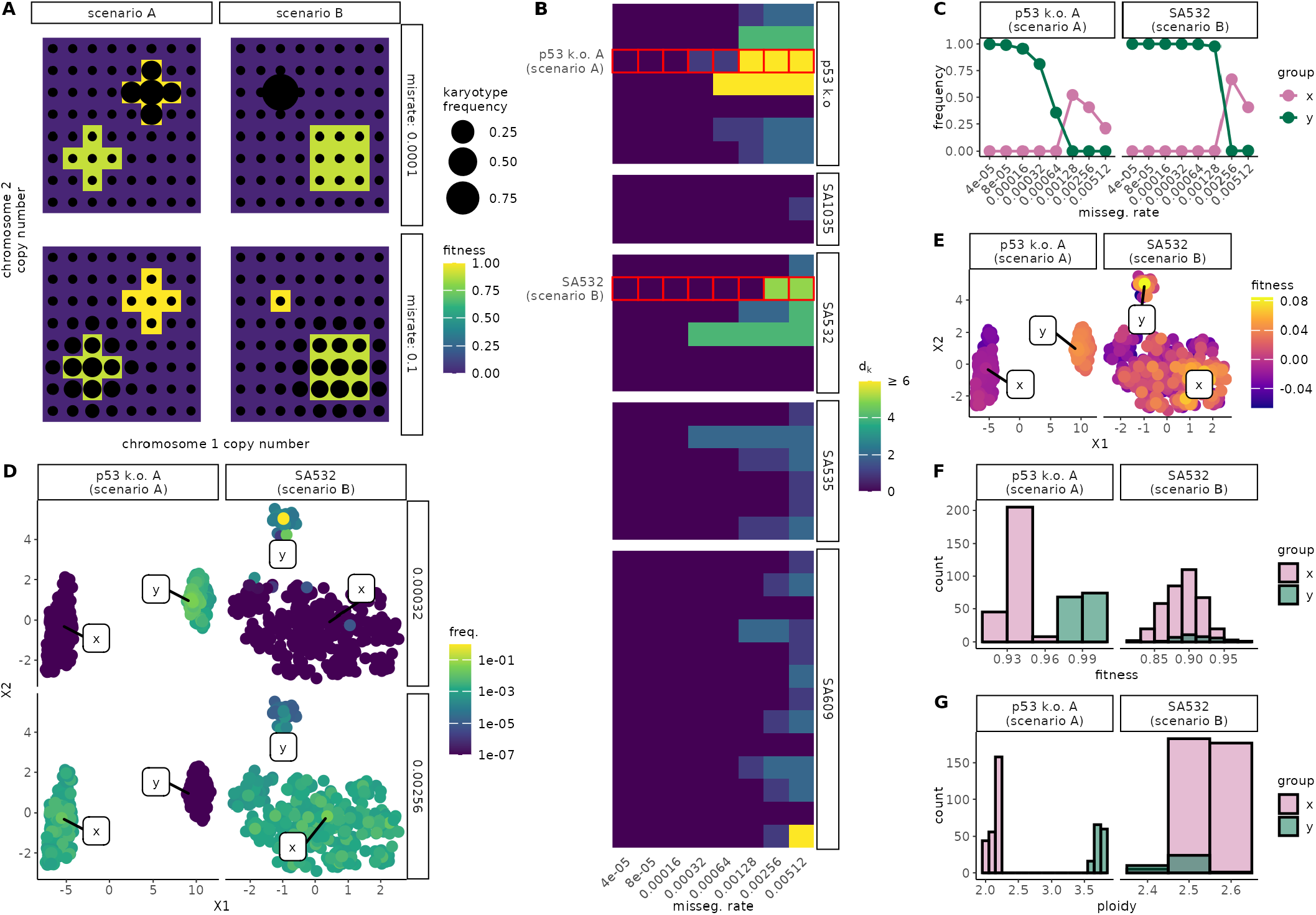
Influence of missegregation rate on karyotype selection. **A)** Hypothetical fitness landscapes for two scenarios (A: ploidy-driven; B: neighborhood-size–driven switches). **B)** Heatmap illustrating screen results: fill color represents the Manhattan distance between the dominant karyotype at steadystate for the lowest missegregation rate compared to the dominant karyotype at the missegregation rate indicated. **C)** Dominant group frequencies (x=high-rate; y=low-rate) in ABM simulations for p53 k.o. A and SA532. **D)** UMAP of all karyotypes colored by steady-state frequency at low (top) vs high (bottom) missegregation rates. **E)** Same UMAPs colored by ALFA-K–estimated fitness. **F**,**G)** Histograms of fitness and ploidy (G) for the x/y groups identified in (C).

To identify rate-dependent switches, we built approximate transition matrices over the charted region of each ALFA-K–fitted landscape and determined the steady state distribution of karyotypes across missegregation rates (Supplementary Section 1.6.1). As an initial screen we searched for landscapes in which the most frequent karyotype (used as a proxy for its quasispecies) differed across missegregation rates. 28 out of 35 tested sublineages met this criteria (Fig. 6B). Two examples were explored in more depth. For both p53 k.o. A and SA532, we retained the 400 most abundant karyotypes and assigned them group membership: group “x” those with abundance negatively correlated to missegregation rate, and group “y” those with abundance positively correlated to missegregation rate. The aggregate frequency of each group shows a clear critical threshold whereupon group dominance within the population flips (Fig. 6C). Moreover, UMAP projections in karyotype space show clear separation between the groups, supporting their interpretation as quasispecies (Figs. 6D–E). Histograms of fitness (Fig. 6F) reveal that in both examples the low-rate dominant group contains the fittest karyotypes. For p53 k.o. A, the high-rate dominants have lower ploidy, whereas for SA532 the selected x/y groups have similar ploidy; by contrast the high-rate dominants enjoy a neighborhood size advantage (Fig. 6G) - supporting the assignment of each cell line to the example scenarios (Fig. 6G). These results demonstrate that missegregation rate can interact with both karyotype fitness landscape shape and ploidy to determine clonal dominance in culture.

## Discussion

To our knowledge, ALFA-K is the first attempt to characterize local regions of karyotype fitness landscapes. We and others have previously modeled karyotypic evolution through the process of missegregation, employing various assumptions about the fitness associated with specific karyotypes. These assumptions range from considering all karyotypes equally fit[19], associating fitness with the density of driver or suppressor genes on each chromosome[20, 21], to correlating fitness negatively with deviations from a euploid state[26]. These models, however, have typically neglected the context in which karyotypic fitness is embedded—factors such as genetic background, tumor microenvironment, and immune interactions profoundly shape the fitness landscape. The vast number of possible karyotypes further complicates the reconstruction of fitness landscapes. Our mathematical model introduces the flexibility needed to begin reconstructing adaptive fitness landscapes, allowing us to extrapolate the fitness of thousands of karyotypes based on the dynamics of just a few subclones.

Whilst the fitness landscapes reconstructed by ALFA-K are not mechanistic, they can be leveraged to provide mechanistic insights into the evolutionary dynamics of the underlying cell populations. It has been observed that cells undergoing whole genome doubling (WGD) exhibit a higher rate of chromosomal alterations than non-WGD cells[27]. The reasons behind this accelerated evolution remain unclear: it could be due to either an increased rate of CNA generation or a heightened tolerance to the deleterious effects of CNAs in WGD cells. We explored this question using an empirical dataset from a cell line that underwent WGD during passaging[23]. Our analysis of the fitness landscape in this line suggests that WGD karyotypes display fewer deleterious CNAs. This supports the notion that WGD may facilitate chromosomal evolution by enhancing cellular robustness to CNAs.

We assessed whether CNA fitness effects generalize across different genomic and experimental contexts by comparing the similarity of Δ*f* profiles between karyotype pairs. Our baseline reference (pairs with substantially different karyotypes from different cell lines) had a Pearson coefficient not significantly above zero. Similarity was significantly higher for karyotypes drawn from the same inferred fitness landscape, which may reflect shared epistatic structure, but could also result from artefactual similarity introduced by using a single model: the Kriging-based fit encourages smoothness across nearby karyotypes, and shared input data can propagate systematic biases to all karyotypes within the same landscape. However, this effect was not limited to shared landscapes—karyotypes differing by fewer than three CNAs showed significantly more similar profiles even after controlling for landscape membership and irrespective of treatment status. This result suggests that modest karyotypic similarity is sufficient to produce convergent CNA fitness effects across otherwise distinct contexts, but that this relationship is fragile—diminishing rapidly as chromosomal distance increases. Whilst a recent analysis found significant correlations in CNA fitness across a large sample of cancers [8], it should be noted that CNAs in that study were all calculated relative to a euploid reference. Our results suggest such analyses could be refined by considering the impact of parental karyotype on CNA fitness effects.

Cells with similar karyotypes inhabiting the same fitness landscape peak can be conceptualized as a “quasispecies”[25]. A prediction of the quasispecies equation is that when mutation is sufficiently rapid, quasispecies will be unable to remain on narrow peaks in the fitness landscape, a concept known as the error threshold [25]. In a novel application of ALFA-K, we investigated the impact of CIN on chromosomal evolution by increasing CIN rates until quasispecies were no longer sustainable on the highest fitness peaks of the landscape. This approach highlighted selection pressures against karyotype quasispecies residing on narrow fitness peaks or in high ploidy regions of the landscape. These findings emerged at missegregation rates of 5-20% per cell division, a range considered plausible for CIN cells[28]. Given that CIN is therapeutically modifiable, our results underscore the potential of using evolutionary principles to steer tumor evolution for long-term control[29]. The ploidy-dependent mechanism builds on prior models, which generally assume equal missegregation rates per chromosome, causing high ploidy karyotypes to missegregate more frequently overall. While this assumption aligns with observations that high ploidy cells missegregate more frequently[27, 30], it warrants further scrutiny. Future models could benefit from integrating metrics such as interferon gamma as a measure of missegregation rate[26], as chromosome missegregations can trigger micronuclei formation, whose rupture activates the cGAS–STING pathway and induces interferon-stimulated gene expression [31]. This could help discern whether a permissive fitness landscape or a propensity to missegregate underlies the relationship between WGD and aneuploidy. One notable limitation of our study is the scarcity of longitudinal data available for training and validating ALFA-K. To mitigate this, we relied heavily on synthetic data and developed metrics to assess the accuracy of the inferred landscapes. However, the absence of independent biological replicates restricted our ability to fully evaluate the quality of our model predictions, as the inherent variability in the system was not well characterized. Future studies with identical biological replicates are essential to further validate ALFA-K[24]. Additionally, data from evolving cell populations inherently focuses on high-fitness regions, making estimation of the fitness impact of deleterious CNAs challenging. ALFA-K attempts to estimate fitness for unobserved karyotypes, using their absence to infer upper fitness limits. However, our estimates for the fitness impacts of deleterious CNAs are likely biased upwards and should be interpreted with caution. Unbiased screens that can accurately estimate the fitness impact of deleterious CNAs would significantly enhance our understanding[27]. A further limitation is that our current implementation treats each chromosome homogeneously, ignoring which allele is lost or gained. In cases where loss of a specific allele has distinct fitness consequences, allele-specific karyotypes would be more appropriate. ALFA-K could be applied directly to such data—simply by tracking each chromosome copy separately—but this would double the number of variables and likely necessitate additional longitudinal data for reliable interpolation.

By quantifying Karyotype Fitness Landscapes (KFL)s, the ALFA-K framework holds significant promise for forecasting long-term evolutionary dynamics in cancer. The application of ALFA-K to hematological malignancies offers a promising future direction for both understanding and treating these cancers. Hematological malignancies, of which an estimated 50% are aneuploid [2], are particularly suited for longitudinal sampling [32], providing the necessary data for accurate fitness landscape reconstruction. Metastatic colonization requires simultaneous adaptation across potentially hundreds to thousands of genes [33, 34], a challenge inefficiently met by sequential point mutations alone. Chromosomal missegregation is a uniquely potent mechanism capable of supplying this complexity in a single leap, yet the multi-generational process of selection on this variation leaves a detectable evolutionary trail. By quantifying the steepness of local slopes and the width of neighboring peaks, ALFA-K converts those trails into in situ maps of evolutionary distance and direction. Similarly, long-term drug responses depend critically on how a tumor population traverses its fitness landscape. Therapeutic interventions that increase chromosomal missegregation rates can push populations beyond their error threshold, causing rapid extinction in rugged KFLs or enabling resistance emergence in smoother ones. In principle, the evolutionary path a clone must travel—from a low-fitness valley to a tissue-specific peak, or from a narrow, drug-sensitive summit to a broader, tolerant one—can be modeled using these inferred landscapes. Thus, ALFA-K charts the terrain that governs these adaptations, providing a quantitative basis to test hypotheses about their timelines without first needing to observe them. Because evolution across KFLs unfolds gradually—over tens to hundreds of cell generations [24, 23]—ALFA-K inference can potentially forecast clinical timelines with actionable lead times, guiding treatment decisions aimed at preventing metastatic outgrowth or the evolution of drug resistance.

## Methods

### The ALFA-K Method for Inferring Karyotype Fitness

#### Overview of the ALFA-K Inference Workflow

We developed ALFA-K (Adaptive Local Fitness landscape for Aneuploid Karyotypes), a method to infer karyotype-specific fitness landscapes and forecast evolutionary trajectories from longitudinal karyotype count data. The core workflow involves three main inference stages (items 1–3) followed by validation (4) and forecasting (5):

1. **Frequent Karyotype Fitness Estimation:** We identify frequently observed karyotypes and estimate their fitness using replicator dynamics.
2. **Single-Step Neighbor Fitness Estimation:** We extend fitness estimates to karyotypes that are one missegregation event away from the frequent karyotypes.
3. **Global Landscape Inference via Kriging:** We use Gaussian process regression (Kriging) to interpolate fitness across the broader karyotype space.
4. **Internal Consistency Check:** We employ a cross-validation procedure to assess the reliability of the inferred landscape.
5. **Evolutionary Forecasting:** We utilize the inferred fitness landscape within a simulation framework to predict future population dynamics.

#### Fitness Estimation for Frequent Karyotypes and their one-MS-step neighbors

The fitness of frequently observed karyotypes (indexed by the set *S*) was modeled using the continuous-time replicator equation, which describes how the relative frequency *x*_*i*_ of a karyotype *i* changes over time under the influence of its fitness *f*_*i*_ and the mean population fitness:

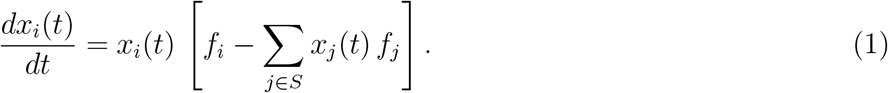

Fitness parameters were first initialized via Quadratic Programming (Supplementary Section 1.2.2) and then refined by maximizing the multinomial likelihood of the observed counts. The detailed mathematical derivations, along with procedures for growth offset correction and bootstrapping, are provided in Supplementary Section 1.2. These fitness estimates for frequent karyotypes were then extended to sparsely observed ‘single-step neighbor’ karyotypes by modeling the mutational flux from their frequent parents:

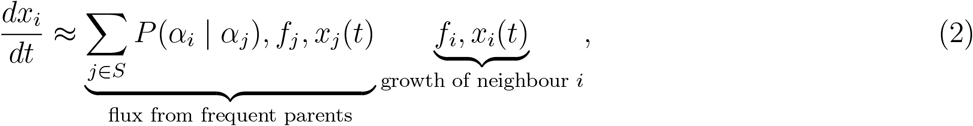

Here *P* (*α*_*i*_ | *α*_*j*_) = Pr **one missegregation with per-chromosome rate** *p* **converts** *α*_*j*_ → *α*_*i*_. The neighbor-specific fitness *f*_*i*_ was inferred by maximizing the likelihood in Eq. S10, which combines the binomial observation term with a Normal prior on the fitness difference, 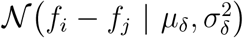 for each parent *j*. The full derivation is provided in Supplementary Section 1.3.

#### Global Landscape Interpolation

A comprehensive fitness landscape was constructed using Gaussian process regression (Kriging) with a Matern kernel (*ν* = 1.5), implemented using the Krig function in the R package fields. This process used the fitness estimates of frequent karyotypes and their one-MS-step neighbors as anchor points. The reliability of interpolated values was assessed via a bootstrap procedure detailed in Supplementary Section 1.4.1.

### Agent-Based Model for Validation and Forecasting

We developed an agent-based model (ABM) to generate synthetic data for validation and to perform forward predictions. In the ABM, cells divide at a karyotype-specific rate (fitness), with a defined per-chromosome missegregation probability (*p*) generating variation. The model was run in two modes: 1) using a known Gaussian-random-field (GRF) fitness landscape to generate synthetic data for benchmarking ALFA-K, and 2) using a look-up table (LUT) of fitness values inferred by ALFA-K to forecast future population dynamics. To emulate cell culture experiments, population size was controlled via simulated serial passaging at a defined maximum capacity (*N*_max_). Full implementation details and simulation parameters are provided in Supplementary Section 2 and Table S1.

## Method Validation

### Validation on Synthetic Data

We ran ABM simulations for 300 days on GRF landscapes with varying complexity (*λ*). Key parameters are listed in Table S1. For testing ALFA-K, we sampled data (simulating experimental measurements) from these simulations. We extracted longitudinal count data for 2, 4, or 8 consecutive passages ending around day 120 of the simulation. This timeframe usually captured populations during active adaptation (Fig. S1E), allowing us to test ALFA-K’s ability to infer ongoing dynamics and predict future evolution. We compared the inferred fitness landscapes against the known ground truth to assess accuracy. The validation showed that inference accuracy, primarily measured by Spearman’s correlation, improved with smoother landscapes and an increased number of sampled time points. The validation procedure and full performance results are detailed in Supplementary Section 3 and Figs. S1-S2.

### Internal Consistency and Forecasting Accuracy

We developed a leave-one-out cross-validation procedure and a resulting metric, CV score, as a key diagnostic for the reliability of the inferred landscape (Supplementary Section 1.5). A positive CV score was strongly correlated with high accuracy against the ground-truth fitness landscape. We then tested the ability of the inferred landscapes to predict future evolution, and found that forecasts from landscapes with a positive CV score were significantly more accurate than random and consistently outperformed simple no-evolution baseline models. Full forecasting methodology and results are available in Supplementary Section 3.4 and Figs. S3-S4.

## Application to Experimental Data

The ALFA-K method was applied to a longitudinal single-cell DNA sequencing dataset [23], which characterized karyotype evolution in immortalized human mammary epithelial cell lines (184-hTERT) and four PDX models. For each cell, we generated a 22-element integer karyotype vector by taking the modal integer copy number across all genomic bins assigned to each autosome (see Supplementary Section 4.1).

## Statistical Analysis

### Analysis of Directional Consistency

To quantify the directional accuracy of evolutionary forecasts in karyotype change, we computed angles between paired karyotype distributions, summarizing each experimental unit by the median of its angles. Each unit corresponded to a serially passaged population derived from one ancestral sample, or in the unrelated control, a pair of such populations from distinct models. The global test statistic *T* was defined as the median of these per-unit medians. To evaluate whether *T* was smaller than expected under random directional change in 22-dimensional space, we performed a Monte Carlo test using the theoretical distribution of angles between random unit vectors. In each of 10,000 iterations, synthetic angles were drawn to match the observed sample sizes per unit, and *T*^∗^ was recomputed. The *p*-value was defined as the proportion of *T*^∗^ values less than or equal to the observed *T* (one-sided for model forecasts and sister passages, two-sided for unrelated controls).

A comprehensive suite of additional metrics was used to evaluate performance; full definitions are provided in Supplementary Section 3.1.

### Prediction of Novel Karyotype Emergence

For each lineage, the latest passage (*S*_0_) was used to predict outcomes in the next (*S*_*t*_). A karyotype was considered <u>novel</u> if it was absent from *S*_0_ but present in *S*_*t*_. Karyotypes were encoded as 22-digit copy-number vectors; Manhattan distance to every *S*_0_ karyotype yielded covariates *d*_1_–*d*_5_, representing the fraction of the *S*_0_ population one to five missegregations away. ALFA-K fitness values (**f**) were computed from landscapes trained only on data up to *S*_0_, ensuring no look-ahead. We retained fits with CV score *>* 0 and at least three passages. When multiple fits terminated at the same passage, we retained only the one with highest CV score, so that each terminal passage defined a single, independent test instance. For each retained fit, we modeled the probability of emergence with a binomial logistic regression: 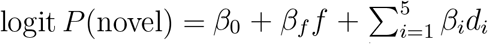. Significance of each predictor was assessed via Wald *z*-tests at *P <* 0.05 (R 4.3, `stats::glm`).

### Analysis of Karyotype Background Influence on Fitness Landscapes

To investigate how the properties of fitness landscapes were shaped by biological context, we first addressed the statistical dependencies caused by overlapping evolutionary trajectories in our data. We implemented a bootstrap framework where, for each of hundreds of replicates, a random set of non-overlapping lineages was algorithmically selected. This method ensured that each replicate of our analysis was based on statistically independent data, allowing for a robust assessment (see Supplementary Section 4.2 for details).

We then used Generalized Linear Mixed Models (GLMMs; `glmmTMB` R package) to analyze the distributions of potential fitness effects (Δ*f*) on these trajectories. We fitted separate models to test how context (PDX vs. in vitro) and drug treatment influenced the magnitude and the variance of fitness effects. These models included nested random effects for the cell line, specific trajectory, and focal karyotype to account for the data structure.

Finally, to assess landscape smoothness, we used a Linear Mixed Model (LMM; `lme4` R package) to model how the similarity between the fitness vectors of two karyotypes (Fisher-z transformed Pearson correlation) decayed as a function of the karyotypic distance between them. The full mathematical specifications for the trajectory selection algorithm and all mixed-effects models are provided in the Supplementary Section 4.2.

### Statistical Modeling of Whole-Genome Doubling (WGD) Effects

To investigate the impact of WGD, karyotypes were first classified as WGD^+^ if their modal chromosome copy number was ≥ 3. We then modeled the dynamics of aneuploidy accumulation over time using non-linear least squares (nls function in R). The average number of altered chromosomes per cell was fitted to an exponential saturation curve where the rate and asymptote parameters were modeled with fixed effects for WGD status and experimental trajectory. The full specification for this non-linear model is detailed in Supplementary Section 4.2.

To directly compare the fitness consequences of mutations, the distributions of single-chromosome gain/loss fitness effects (Δ*f*) for WGD^+^ versus WGD^−^ states were compared using a permutation Kolmogorov-Smirnov (KS) test. To avoid pseudo-replication, mean effects were calculated per WGD status within each landscape fit before comparison. Significance was then assessed against 10^4^ permutations of the WGD labels among the landscape fits.

## Data Availability

ALFA-K is available here: https://github.com/Richard-Beck/alfakR. Scripts and data needed to reproduce analysis presented in this work are available here: https://github.com/Richard-Beck/ALFA-K.

## Acknowledgements

We are grateful to the members of the Integrated Mathematical Oncology (IMO) department at Moffitt for their valuable feedback on this manuscript. The insightful comments and suggestions provided during numerous department meetings were instrumental in strengthening the final work.

## Supplementary Information

### 1 The ALFA-K Method: Inferring and Forecasting Karyotype Evolution

Here, we provide the full details of our ALFA-K (Adaptive Local Fitness landscape for Aneuploid Karyotypes) inference method. ALFA-K infers karyotype fitness from longitudinal count data and uses this information to forecast evolutionary trajectories. Central to our approach is the concept of fitness, which we interpret as the intrinsic net growth rate associated with each distinct karyotype. We assume that the observed karyotype counts obtained at different time points are the result of uniform sampling from the underlying population. This sampling process links the observed data to the unobserved, true relative frequencies of karyotypes within the population.

#### 1.1 Overview of the ALFA-K Workflow

ALFA-K takes as input a longitudinal count matrix *Y*, whose entry *y*_*it*_ is the count of karyotype *i* at time *t*. The core workflow involves three main inference stages (items 1–3) followed by validation (4) and forecasting (5):

1. **Frequent Karyotype Fitness Estimation:** Identify frequently observed karyotypes and estimate their fitness using replicator dynamics.
2. **Single-Step Neighbor Fitness Estimation:** Extend fitness estimates to karyotypes that are one missegregation event away from frequent karyotypes.
3. **Global Landscape Inference via Kriging:** Use Gaussian process regression (Kriging) to interpolate fitness across the broader karyotype space, leveraging the estimates from frequent clones and their neighbors.
4. **Internal Consistency Check:** Employ a cross-validation procedure to assess the reliability and generalization capability of the inferred landscape.
5. **Evolutionary Forecasting:** Utilize the inferred fitness landscape within a simulation framework to predict future population dynamics.

Each stage is detailed below.

#### 1.2 Step 1: Fitness Estimation for Frequent Karyotypes

We begin by focusing on inferring the fitness of frequently observed karyotypes. A key simplification in this step is modeling these types as effectively present from the beginning of the observation window, each with an initial frequency estimated directly from the data alongside its fitness (*f*_*i*_). This approach circumvents the need to reconstruct potentially complex and underdetermined evolutionary pathways leading to these frequent states, while still accommodating clones that may have emerged later (captured by a small fitted initial abundance). Further simplifying assumptions for this step include neglecting stochastic extinction (acceptable due the inference’s low sensitivity to the distinction between rare and absent states), mutations <u>between</u> these frequent types, density-dependent growth, and frequency-dependent selection. By making these targeted simplifications, we obtain robust, data-driven fitness estimates for the most prevalent karyotypes, which serve as a critical foundation for the subsequent steps of the pipeline.

##### 1.2.1 Definition and Model

A karyotype *i* is deemed frequent if its total count across all time points satisfies ∑_*t*_ *y*_*it*_ ≥ *N*, for a user-defined threshold *N*. Let *S* be the set indexing these frequent karyotypes. We model the dynamics of each frequent karyotype *i* ∈ *S* using the continuous-time replicator equation, which describes frequency changes under selection:

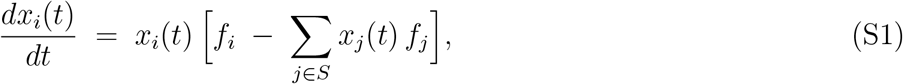

where *x*_*i*_(*t*) is the frequency of karyotype *i* at time *t* relative to other frequent karyotypes in *S*, and *f*_*i*_ is its constant, intrinsic fitness parameter (net growth rate).

##### Interpretation (Replicator vs. Exponential Growth)

The replicator equation emerges naturally from assuming independent exponential growth for each karyotype. If the absolute abundance *y*_*i*_(*t*) of karyotype *i* follows 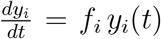 (leading to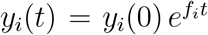), then its relative frequency 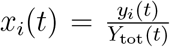, where *Y*_tot_(*t*) = *#x2211;*_*j*_ *y*_*j*_(*t*), indeed satisfies Equation S1. Thus, inferring *f*_*i*_ under the replicator model is equivalent to inferring the exponential growth rate from relative frequency data.

##### 1.2.2 Fitness Inference Procedure

We estimate the fitness vector **f** = {*f*_*i*_}_*i* ∈*S*_ in two steps:

##### Initial Estimation via Quadratic Programming (QP)

To obtain initial fitness estimates, we linearize the replicator dynamics over short time intervals Δ*t*. Let *x*_*it*_ = *y*_*it*_*/N*_*t*_, where *N*_*t*_ = ∑_*j*∈*S*_ *y*_*jt*_ is the total count of frequent karyotypes at time *t*. The discrete frequency change is Δ*x*_*i,t*_ = (*x*_*i,t*+1_ − *x*_*i,t*_)*/*Δ*t*.

In vector form, Equation S1 can be written as 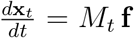, where 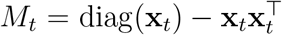. Approximating the derivative gives Δ**x**_*t*_ ≈ *M*_*t*_**f**. We find the initial **f** by minimizing the sum of squared errors across all time intervals:

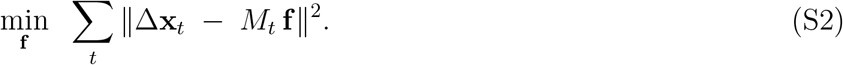

T his least-squares problem can be formulated as a quadratic program. Defining 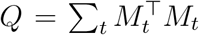 and 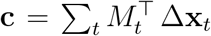, the objective becomes minimizing 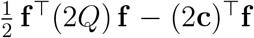. We add a small regularization term *ϵI* for numerical stability and impose the constraint _∑*i*∈*S*_ *f*_*i*_ = 0 to set the mean fitness to zero, yielding the QP:

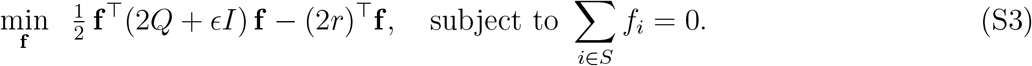

##### Refinement via Joint Likelihood Optimization

We refine the initial estimates {*f*_*i*_} by maximizing the likelihood of the observed counts *Y* = {*y*_*i,t*_} under the full replicator model. The analytical solution to Equation S1 for constant fitness is:

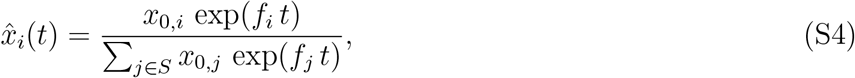

where {*x*_0,*i*_} are the initial frequencies at *t* = 0 (∑_*i*∈*S*_ *x*_0,*i*_ = 1). Assuming the observed counts *Y*_*t*_ ={*y*_*i,t*_} at time *t* follow a multinomial distribution with *N*_*t*_ total trials (cells) and probabilities 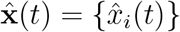, the likelihood is:

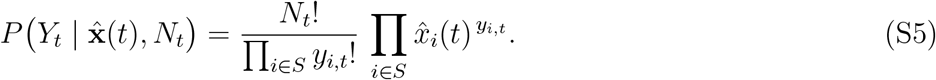

We maximize the total log-likelihood 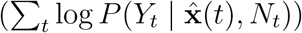 jointly over the fitness parameters {*f*_*i*_} and the initial frequencies {*x*_0,*i*_}, subject to _∑*i*∈*S*_ *f*_*i*_ = 0. The QP solution provides the starting values for the optimization.

##### Growth Offset Correction for Passaging Experiments

In cell culture experiments involving passaging, the overall population growth rate can be estimated. If *n*_0_ and *n*_*b*_ are the total cell counts at the start and end of a passage of duration Δ*t*, the observed mean growth rate is *g*_0_ = ln(*n*_*b*_*/n*_0_)*/*Δ*t*. We shift the infer red relative fitness values {*f*_*i*_} by a constant so that the mean fitness predicted by the replicator model (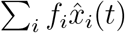, averaged over time) matches the observed mean growth rate *g*_0_.

##### Bootstrapping

To estimate uncertainty in fitness parameters, both the QP and maximum-likelihood fitting steps are repeated across multiple bootstrap resamples of the input count data {*y*_*it*_}. The output of this step is a distribution of fitness estimates per each frequent karyotype.

### 1.3 Step 2: Fitness Estimation for one-MS-step neighbors

In this step, we extend fitness estimates to the sparsely observed karyotypes that are one gain or loss event (‘one-MS-step neighbors’) away from the frequent karyotypes identified in Step 1. Due to their low counts, full dynamic modeling is impractical. Instead, we approximate their fitness (*f*_*i*_) based on the inferred mutational flux from their frequent parents (*j* ∈ *S*). This approach assumes that these neighbors primarily arise through such single missegregation events (governed by a missegregation probability *P* (*α*_*i*_ |*α*_*j*_) dependent on the per-chromosome rate *p*); that flux from rare-to-rare or rare-to-frequent types is negligible; and that the missegregation rate *p* is constant and known (or estimated). Let:

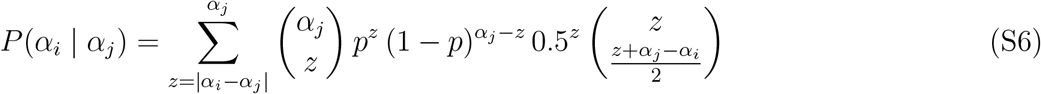

be the probability of a missegregation event producing *i* (copy number *α*_*i*_) from *j* (copy number *α*_*j*_), where *p* is the per-chromosome missegregation rate [26, 19].

#### 1.3.1 Fitness Inference via Flux Approximation and Likelihood

The rate of change of neighbor *i*’s frequency *x*_*i*_(*t*) is approximated as:

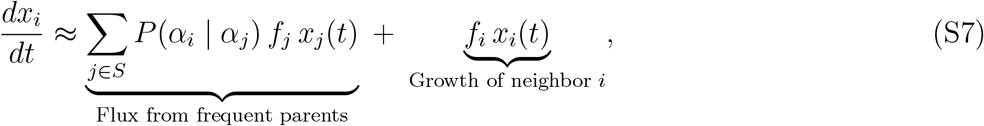

neglecting missegregation from other rare types and back-mutation to frequent types. This ODE can be solved approximately. If we consider flux from a single parent *j* starting at effective time *t*_birth,*j*_ and approximate *x*_*j*_(*t*) ≈ *x*_*j*_(*t*_birth,*j*_) exp(*f*_*j*_ [*t* − *t*_birth,*j*_]), the contribution 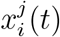 from parent *j* evolves according to:

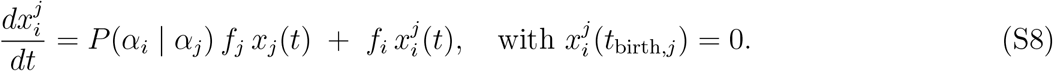

The solution relates the ratio 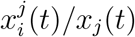 to the fitness difference (*f*_*i*_ − *f*_*j*_):

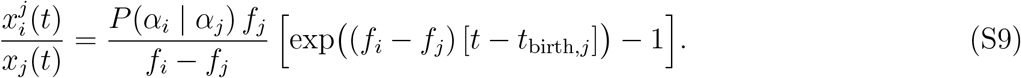

The total expected frequency is 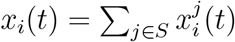, under the approximation that *x*_*i*_ remains small.

We estimate *f*_*i*_ for neighbor *i* by maximizing a likelihood function that combines the probability of observing its counts *y*_*i,t*_ with a prior on fitness differences relative to its parents. Let 𝒫 (*i*) ⊆ *S* be the set of frequent parents of *i*. We assume fitness differences *δ*_*ij*_ = *f*_*i*_ − *f*_*j*_ follow a Gaussian distribution with mean *µ*_*δ*_ and standard deviation *σ*_*δ*_ (estimated empirically from all neighbor-parent pairs). The likelihood for *f*_*i*_ is:

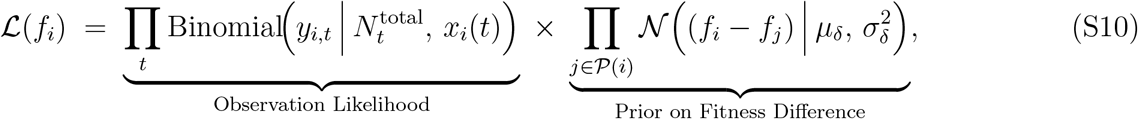

where 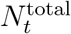 is the total cell count at time *t*, and *x*_*i*_(*t*) depends on *f*_*i*_ via Equation S9. We maximize log ℒ (*f*_*i*_) with respect to *f*_*i*_.

The single-step neighbor fitnesses are estimated within the same boostrapping loop as the frequent karyotypes, using the same resampled data. Thus the output of this step is a distribution of fitness estimates per each neighbor karyotype.

### 1.4 Step 3: Fitness Inference for All Other Karyotypes via Kriging

To build a more comprehensive fitness landscape, this step estimates fitness for remaining viable karyotypes (those neither frequent nor direct one-MS-step neighbors of frequent types). We employ Gaussian process regression (Kriging) to interpolate fitness values across the karyotype space. This interpolation uses the fitness estimates derived in Steps 1 and 2 as anchor points. The core assumption underpinning this approach is that fitness constitutes a relatively smooth function over the space of karyotypes (represented by copy number vectors), and specifically that the chosen Matern kernel (with *ν* = 1.5) adequately captures this smoothness structure. Karyotypes containing zero copies of any chromosome are deemed non-viable and excluded prior to interpolation. It is important to note that, as with any interpolation method, the reliability of Kriging predictions diminishes significantly for karyotypes distant from the anchor points in the high-dimensional space. Therefore, whilst in principle we could apply this interpolation across the entire karyotype space, in practice we limit our predictions to single-step neighbours of karyotypes with fitness estiamtes from the prior two steps. Kriging was implemented using the `Krig` function in the R package “fields”, with all parameters except the aforementioned kernel set at default values. The default setting (*ν* = 1.5) in `fields::Krig()` corresponds to an exponential kernel, which assumes the fitness landscape is maximally rough. This may understate the degree of local structure in karyotype space, where similar copy-number profiles often exhibit related fitness. Increasing *ν* to 1.5 allows for limited smoothness while preserving flexibility, in the absence of definitive information about the landscape’s regularity.

### 1.4.1 Bootstrap Procedure for Uncertainty and Decorrelation

The Kriging step is performed within a separate bootstrap procedure. At every bootstrap iteration we keep the karyotype coordinates fixed but resample, with replacement, the anchor fitness estimates generated in Steps 1–2. A fresh Gaussian-process model is fitted to each resampled dataset and used to predict fitness over the full evaluation set of karyotypes. This helps in obtaining uncertainty estimates for the Kriging predictions and is specifically designed to reduce potential correlations between the errors in fitness estimates for frequent clones and the errors for their neighbors (which depend on the parent’s fitness). By using a sufficient number of bootstrap iterations, we obtain a distribution of fitness predictions for each interpolated karyotype. We summarize this distribution by its mean and standard deviation, assuming normality, providing a compact representation of the inferred landscape and its uncertainty.

## 1.5 Assessing Internal Consistency: Cross-Validation Procedure

To evaluate the internal consistency and predictive reliability of the inferred fitness landscape, particularly in the absence of ground truth experimental data, we employ a cross-validation procedure. This procedure assesses the extent to which the fitness landscape exhibits local structure, specifically testing whether the fitness of any given karyotype and its immediate neighbors can be reasonably predicted from the landscape inferred using data from other related karyotypes. A high degree of predictability indicates good generalization capability and robustness of the overall fitness landscape constructed in the preceding steps, while a low degree suggests otherwise.

### 1.5.1 Method

The procedure focuses on the frequent karyotypes and their neighbors: For each frequent karyotype *i* ∈ *S*:

1. Temporarily remove frequent karyotype *i* and all of its one-MS-step neighbors from the set of known fitness values (obtained in Sections 1.2 and 1.3).
2. Re-fit the Kriging model (Section 1.4) using the fitness data from the remaining frequent karyotypes and their neighbors.
3. Use this re-fitted Kriging model to predict the fitness values for the held-out karyotype *i* and its neighbors. Let these be the cross-validated predictions *g*_*c*_.

This process is repeated for every frequent karyotype *i* ∈ *S*. Let 𝒞 be the set of all karyotypes whose fitness was predicted during this CV procedure (i.e., all frequent karyotypes and all their one-MS-step neighbors). We compare the cross-validated predictions *g*_*c*_ for *c* ∈ 𝒞 with the original estimates *f*_*c*_ obtained using the full dataset.

### 1.5.2 Cross-Validation Metric 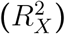

We quantify the agreement between the original estimates and the cross-validated predictions using a rescaled coefficient of determination, 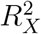 :

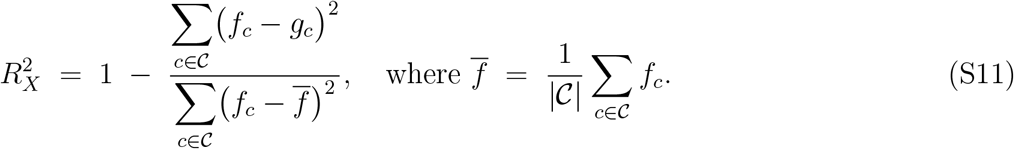

A high 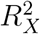 suggests that the fitness estimates for individual karyotypes are well-supported by the surrounding fitness landscape inferred from other related clones, indicating internal consistency and good generalization. A low or negative 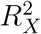 suggests potential issues like overfitting to noise in the frequency data of the held-out clone or poor interpolation by the Kriging model.

### 1.6 Predicting Future Evolution: Forecasting Simulation

ALFA-K can use the inferred fitness landscape (represented by mean fitness 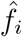 and standard deviation *σ*_*i*_ for each karyotype *i*) to forecast future population dynamics. To predict dynamics of evolving populations, we used an agent-based model (Section 2).

#### 1.6.1 Calculating the Steady-State Distribution

We determined the steady-state relative frequency distribution of co-existing karyotypes (**x**_ss_) under constant conditions by analyzing the linear system governing the dynamics of absolute cell abundances (**y**) as described previously [26]. This system is described by the ordinary differential equation 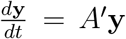, where the matrix *A*^*′*^ incorporates both fitness-dependent growth and mutation rates derived from the per-division missegregation probabilities (*Q*). Specifically, the off-diagonal elements 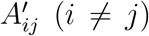 represent the rate at which type *j* produces type *i* via missegregation, given by the product of the parent fitness and the per-division probability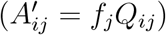. The diagonal elements 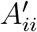 represent the net growth rate of type *i*, accounting for its intrinsic fitness *f*_*i*_ and its loss rate due to mutating into other types 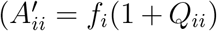, where *Q*_*ii*_ = −∑_*k* ≠ *i*_ *Q*_*ki*_ is derived from the per-division probabilities). We numerically constructed this matrix *A*^*′*^ using the inferred fitnesses **f** and the mutation probabilities *Q* derived from the per-chromosome rate *p*. The steady-state relative frequency distribution **x**_ss_ corresponds to the normalized dominant eigenvector **v**_*max*_ of *A*^*′*^. This vector **v**_*max*_, associated with the eigenvalue of *A*^*′*^ having the largest real part (which dictates the long-term asymptotic growth rate of the system), was computed efficiently using sparse matrix eigensolvers. Finally, the eigenvector was normalized such that its elements sum to one (**x**_ss_ = **v**_*max*_*/* ∑ _*k*_ *v*_*k,max*_) to yield the probability distribution representing the steady state.

## 2 Agent-Based Model (ABM)

### 2.1 Overview

We simulate the evolution of chromosome-level copy-number profiles in a population of cultured cells assumed to be <u>well mixed</u>, meaning that each cell experiences the same environment and competes equally with every other cell; no spatial structure is modelled. The simulator supports two use cases, corresponding to separate analyses in our manuscript:

**Synthetic-data generation (GRF mode)**: Fitness is defined by a known Gaussian-random-field (GRF) function, allowing us to benchmark how accurately ALFA-K recovers the underlying landscape. **Forward prediction (LUT mode)**: Fitness values inferred by ALFA-K are supplied as a look-up table, and the simulator predicts future karyotype dynamics under those selective pressures.

Cells can only change in number or genotype through mitotic division. Division rates are karyotype specific; chromosomal missegregation during division generates variation. Spontaneous death is not explicitly modelled, but cells are removed if they produce inviable daughters (with zero copies of any chromosome class) or during periodic passaging (see 2.1).

#### Software implementation

The simulator is written in C++17 and exposed to R through `Rcpp`. Source code, documentation, and installation instructions are available in the `alfakR` package (https://github.com/Richard-Beck/alfakR).

#### Key model objects

- **Karyotypes** are stored as fixed-length integer vectors, one element per autosome. All copy numbers are strictly positive.
- **Population state** maps each unique karyotype to its current cell count.
- **Fitness specification** is supplied either as a numeric table (LUT mode) or by a Gaussian-random-field (GRF) function,

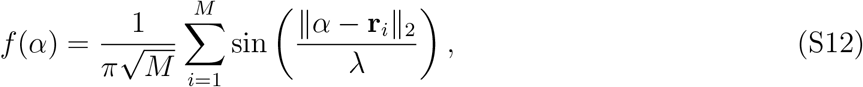

where *α* is the karyotype vector, {**r**_*i*_} are user-defined centroids, and *λ* is the GRF wavelength parameter.

### Stochastic division-and-segregation algorithm

At each discrete time increment Δ*t*, the simulator processes every karyotype present (consistent with previous models [26, 19]) as follows:

1. **Number of divisions**. Let *n*_*i*_ be the number of cells with a given karyotype and *f* its fitness. The expected number of divisions is

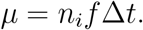

The realised number of divisions is sampled from a Poisson distribution with mean *µ*, but cannot exceed *n*_*i*_.
2. **Segregation outcome**. Let *C* be the total number of chromosomes in the parent cell, and *p* the per-chromosome missegregation probability.
  - With probability (1 − *p*)^*C*^, the division is faithful and both daughters match the parent.
  - Otherwise, the division is error-prone. The number of chromosomes that mis-segregate is sampled as

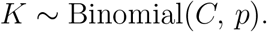
3. **Daughter formation**. Exactly *K* chromosomes are sampled uniformly without replacement from the *C* total. Each mis-segregating chromosome is randomly allocated to one daughter (with probability 1*/*2), resulting in a +1 copy number change in that daughter and −1 in the other. Any daughter with a zero in any chromosome class is discarded.
4. **Population update**. Parent cells that divided are removed; viable daughters are added to the population.

All stochastic sampling uses a Mersenne Twister engine, seeded by the user or from hardware entropy if no seed is provided.

### Population control (serial passaging)

To emulate the dynamics of cell-culture and xenograft experiments—where cells grow to capacity and are then re-seeded at lower density—the simulator imposes a user-defined maximum population size *N*_max_. When the total cell count exceeds this threshold, each karyotype is independently down-sampled via a binomial draw that retains a fixed fraction *s* of its cells. This reduces the total number of cells while preserving relative karyotype abundances.

## 3 Validation of ALFA-K using Synthetic Data

To rigorously evaluate ALFA-K’s performance and understand its limitations, we generated synthetic datasets where the ground truth fitness landscape and evolutionary dynamics are known. This approach allows us to directly compare inferred fitness landscapes against the known ground truth, assess the impact of factors such as landscape complexity, sampling frequency, and noise on inference accuracy, evaluate the reliability of the cross-validation score 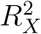 as a diagnostic tool, and test the accuracy of evolutionary forecasts against known future population states. These synthetic datasets were generated using an Agent-Based Model (ABM) to simulate population evolution on Gaussian Random Field (GRF) fitness landscapes.

### 3.0.1 Simulation Setup and Parameters

We ran ABM simulations for 300 days on GRF landscapes with varying complexity (*λ*). Key parameters are listed in Table S1. Populations evolving on complex landscapes (low *λ*) often exhibited punctuated evolutionary dynamics, whereas those on smoother landscapes (high *λ*) showed more gradual fitness increases (Fig. S1A-D).

**Table S1:** Parameters used in ABM simulations. ABM simulations are used: to predict karyotype evolution for empirical datasets (Sim 1); to generate synthetic data to test ALFA-K (Sim 2); and to predict evolution of the synthetic cell populations based on ALFA-K fitness landscapes (Sim 3). {**r**_*i*_} and the fitness table represent optional arguments whose respective presence or absence is indicated with ‘y’ or ‘n’.

| Description | Symbol | Sim 1 | Sim 2 | Sim 3 | Comments |
| --- | --- | --- | --- | --- | --- |
| Per-chromosome missegregation probability | $p$ | | $5 \times 10^{-5}$ | | $(10^{-5}-10^{-1})$ [2, 35, 36] |
| Time increment per step | $\Delta t$ | | 0.1 | | — |
| Number of simulation steps | $\ell$ | variable | 3000 | variable | — |
| Population cap (passaging threshold) | $n_{\max}$ | $10^7$ | $2 \times 10^6$ | $2 \times 10^6$ | Corresponds to small tumour fragments or large cultures [37] |
| Survival fraction after passaging | $s$ | | 0.01 | | Typical value for seeding PDX models [38] |
| Initial population size | $N_0$ | $10^5$ | $5 \times 10^4$ | $2 \times 10^5$ | Typical values for seeding PDX models [39] |
| GRF wavelength | $\lambda$ | — | 0.2–1.6 | — | |
| GRF centroids | $\{\mathbf{r}_i\}$ | y | n | y | — |
| Fitness table (LUT mode only) | — | y | n | y | — |

For testing ALFA-K, we sampled data (simulating experimental measurements) from these simulations. We extracted longitudinal count data for 2, 4, or 8 consecutive passages ending around day 120 of the simulation (Fig. S1E). This timeframe usually captured populations during active adaptation, allowing us to test ALFA-K’s ability to infer ongoing dynamics and predict future evolution.

### 3.1 Metrics for Performance Evaluation

We used several metrics to compare ALFA-K’s inferred fitness landscapes (*f* ^*pred*^) against the ground truth (*f* ^*true*^) from the GRF, and to compare forecasted population states (**x**^*p*^) against the true future states from the ABM (**x**^*a*^).

**Variables for Metrics** Let *f* ^*pred*^ and *f* ^*true*^ be predicted and true fitness vectors. Let **x**^*p*^, **x**^*a*^, and **x**^0^ be predicted, actual, and baseline frequency vectors, respectively. Let **k**_*i*_ be the vector representation of karyotype *i*. The centroid is *c*(**x**) = ∑ _*i*_ **k**_*i*_*x*_*i*_*/ ∑* _*i*_ *x*_*i*_.

**Spearman’s Correlation (*ρ*)** Measures rank correlation between *f* ^*pred*^ and *f* ^*true*^:

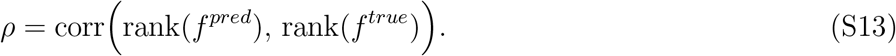

**Figure S1:**
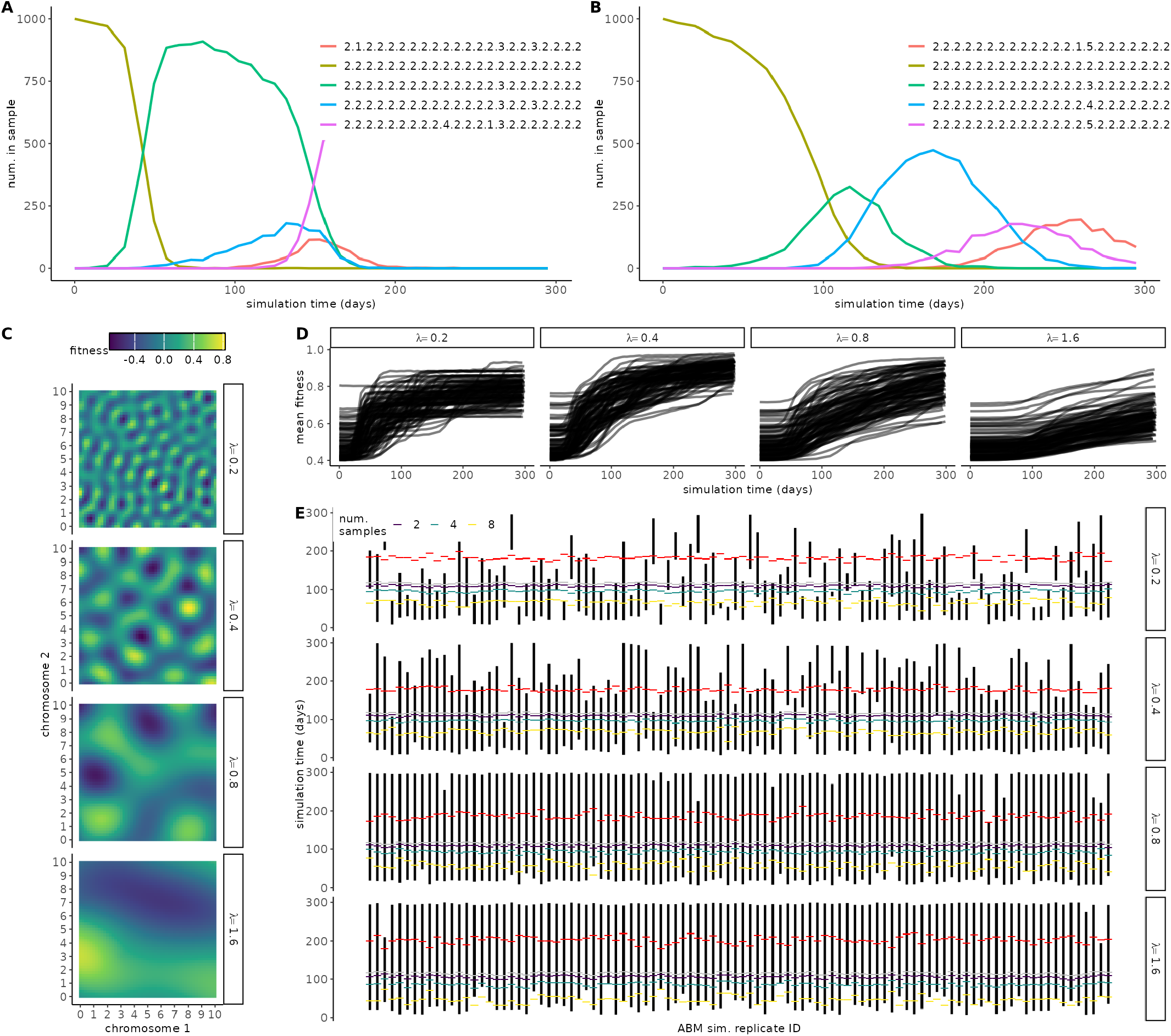
In silico test populations and sampling strategy. (A-B) Example simulation output for ABM cell populations evolving on GRF fitness landscape with A) *λ* = 0.2 or B) *λ* = 1.6. Each coloured line represents the longitudinal frequency of a different karyotype. (C) Increasing the wavelength (*λ*) results in GRF with decreasing complexity. (D) Mean fitness of ABM cell populations evolving on artificial fitness landscapes of varying complexity (as determined by *λ*). (E) Overview of ABM sampling strategy. Black bars represent the longest period of continuous evolutionary progress in each ABM simulation as measured by consecutive passages with fitness increases. Colored points indicate the time intervals included in ALFA-K training for different numbers of timepoints, with grey points representing the latest included timepoints and red points showing the latest predictions.

Pearson’s Correlation (*r*)

Measures linear correlation between *f* ^*pred*^ and *f* ^*true*^:

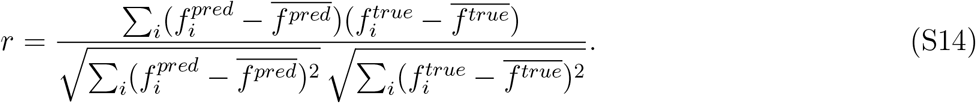

**Rescaled** 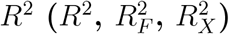

Proportion of variance explained, computed on mean-centered data:

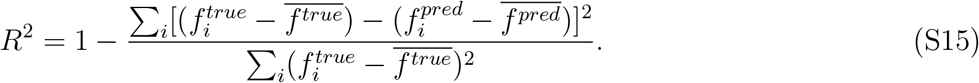

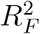 is computed on frequent karyotypes only. 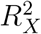 is the cross-validation metric (Eq. S11).

**Angle Metric (*θ*)** Quantifies directional alignment of trajectories via centroids. Displacement vectors: **v**^*p*^ = *c*(**x**^*p*^) − *c*(**x**^0^), **v**^*a*^ = *c*(**x**^*a*^) − *c*(**x**^0^).

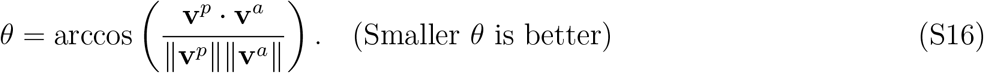

The null distribution CDF for *θ* between two random unit vectors in ℝ^*N*^ is:

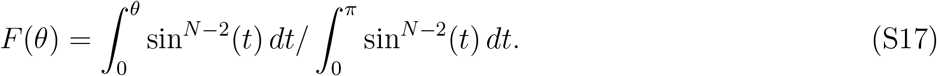

**Cosine Similarity (CS)** Compares frequency profiles **x**^*p*^ and **x**^*a*^:

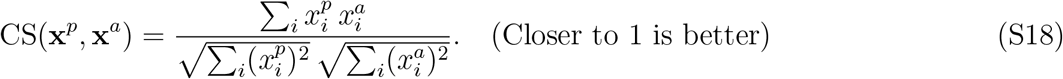

**Euclidean Distance (*d*_*E*_)** Distance between centroids *c*(**x**^*p*^) and *c*(**x**^*a*^):

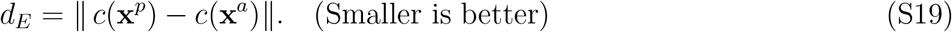

**Overlap Coefficient (Ω)** Fraction of shared population mass:

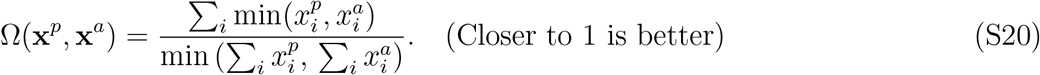

**Wasserstein Distance (*d*_*W*_)** Earth Mover’s Distance between distributions **x**^*p*^ and **x**^*a*^, requiring minimal cost to transform one to the other based on euclidean distance *d*(**k**_1_, **k**_2_) between karyotypes:

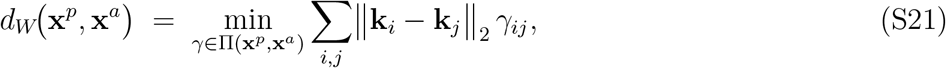

where Π(**x**^*p*^, **x**^*a*^) is the set of transport plans whose row and column sums equal **x**^*p*^ and **x**^*a*^, respectively.

A smaller *d*_*W*_ indicates that the predicted distribution is closer to the actual one.

## 3.2 Evaluating Fitness Landscape Inference Accuracy

We applied ALFA-K to the sampled synthetic data and compared the inferred fitness landscapes to the known GRF ground truth using metrics defined in Section 3.1. Accuracy generally improved with smoother landscapes (higher *λ*), a higher threshold *N* for defining frequent karyotypes, and more sampled time points (Fig. S2A). These factors help mitigate the impact of sampling noise and demographic stochasticity, especially for low-frequency clones.

The analysis revealed that accurate fitness estimation for the initial set of frequent karyotypes is crucial. Fits where the frequent-subset accuracy 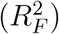 was poor, or where the frequent set was very small, rarely yielded accurate global landscapes (positive global *R*^2^) (Fig. S2B). Errors in the initial frequent set estimation tend to propagate through the neighbor extension and Kriging steps. An example simulation highlights this (Fig. S2C-E). With limited training data (2 passages), stochastic fluctuations in frequency trajectories could lead to incorrect ranking of closely competing clones. Providing more data (8 passages) allowed ALFA-K to resolve the long-term trends and correctly infer the fitness ranking, even with the same *N* threshold.

A Sankey diagram summarizing all fits (Fig. S2F) confirmed that poor global accuracy 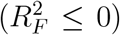 was strongly associated with either poor initial frequent set accuracy (*R*^2^ ≤ 0) or an insufficient number of frequent karyotypes identified, regardless of landscape complexity or number of passages. This underscores the importance of the initial frequent karyotype identification and fitness estimation step.

## 3.3 Evaluating Cross-Validation as a Diagnostic Heuristic

Since ground truth fitness is unknown in real experiments, we evaluated the utility of the cross-validation score 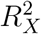 (Section 1.5) as a proxy for inference reliability using our synthetic data. The 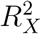 scores calculated on the synthetic datasets showed trends consistent with direct accuracy metrics: scores were lower for more rugged landscapes (low *λ*) and when fewer training passages were used (Fig. S3A). This suggests 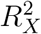 captures aspects related to the difficulty of inference. Importantly, we found a strong correlation between the sign of the cross-validation score and the sign of the true accuracy (*R*^2^) against the ground truth. Fits achieving a positive *R*^2^ were significantly enriched for also having a positive 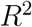 (Fig. S3B). Furthermore, landscapes inferred with 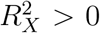 0 showed substantially higher accuracy across all metrics (*ρ, r, R*^2^) compared to those with 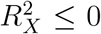 (Fig. S3C). These results support the use of 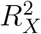 as a diagnostic heuristic. While not a guarantee of accuracy, a positive 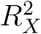 indicates internal consistency and suggests the inferred landscape is more likely to be reliable.

## 3.4 Evaluating Evolutionary Forecasting Performance

Finally, we tested the ability of ALFA-K’s inferred fitness landscapes to predict future evolutionary dynamics in the ABM simulations. We used the forecasting method described in Section 1.6 to simulate evolution forward from the last training time point and compared the predicted population state to the true state from the ABM simulation at later times.

## 3.4.1 Directional Accuracy (Angle Metric)

We assessed directional accuracy using the angle metric *θ* (Section 3.1), where lower values indicate better alignment between predicted and true evolutionary vectors. ALFA-K forecasts consistently achieved better-than-random directional accuracy (ECDFs shifted left from the null distribution, Fig. S4A). Accuracy generally improved with more training samples and decreased slightly for longer prediction horizons (more passages forecast). Crucially, forecasts generated from landscapes with good internal consistency 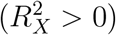 were significantly more accurate directionally than those from landscapes with 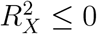.

## 3.4.2 Comparison Against No-Evolution Baselines

We also compared ALFA-K forecasts against simple baseline models that assume no evolution (i.e., predicting the final observed state persists). We measured the fraction of times ALFA-K forecasts were ‘better’ than the baseline according to various metrics (Cosine Similarity, Euclidean Distance, etc.). When trained on sufficient data (e.g., 4+ passages) and achieving a positive cross-validation score 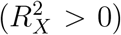, ALFA-K forecasts outperformed these static baselines in roughly 75% of cases across different metrics and forecast horizons (Fig. S4B). In contrast, forecasts from fits with 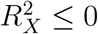 rarely surpassed the baselines.

**Figure S2:**
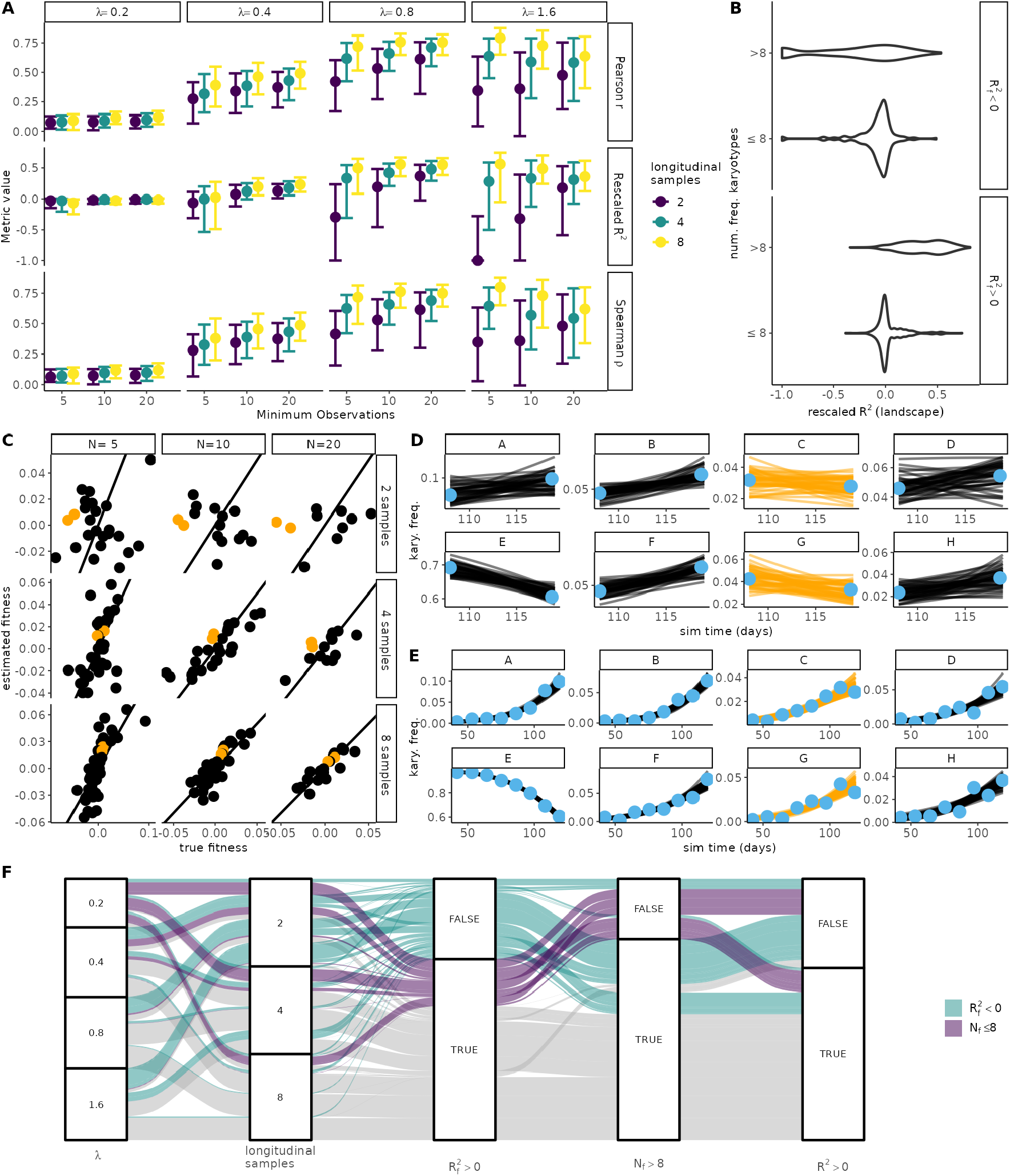
Performance of ALFA-K fitness inference on simulated landscapes. (A) Accuracy metrics (*ρ, r, R*^2^; rows) vs. ground truth GRF fitness, shown as a function of landscape complexity (*λ*, columns), frequent karyotype threshold (*N*, x-axis), and number of sampled time points (colour). Boxes: 10th, 50th, 90th percentiles. (B) Global rescaled *R*^2^ vs. frequent-subset 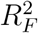 and number of frequent karyotypes identified. (C) Example: Estimated vs. true fitness for one simulation (*λ* = 0.8) with varying hyperparameters. Orange points highlight three clones whose rank order is inverted with limited data. (D–E) Frequency trajectories (observed points, fitted lines) for the top 8 karyotypes from (C) using *N* = 20 and (D) 2 passages or (E) 8 passages. Orange lines correspond to the mis-ranked clones in (C, D). (F) Sankey diagram summarizing accuracy across all fits, stratified by input parameters and intermediate results (sign of 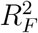, number of frequent karyotypes abbreviated as *N*_*F*_) leading to final global *R*^2^ sign.

## 3.4.3 Conclusion from Forecasting Evaluation

The forecasting results further validate ALFA-K and the utility of the 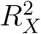 diagnostic. Landscapes deemed reliable by cross-validation not only reflect the known fitness landscape more accurately but also possess significantly better predictive power regarding future evolutionary trajectories compared to unreliable landscapes or simple no-evolution assumptions.

**Figure S3:**
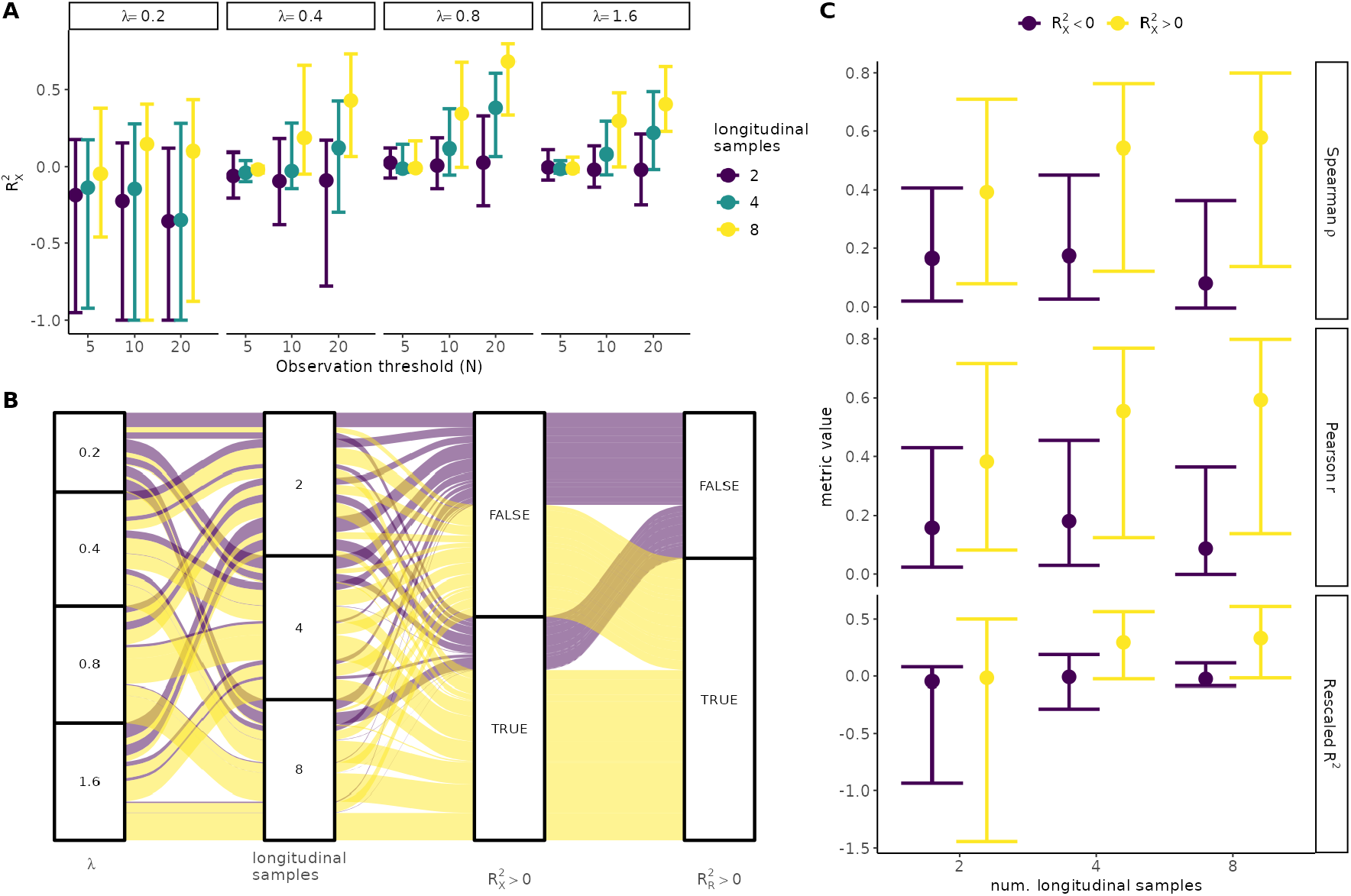
Cross-validation metric 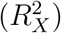 as a diagnostic heuristic for ALFA-K performance on ABM data. (A) Cross-validation scores 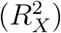 vs. landscape complexity (*λ*, columns), frequent karyotype threshold (*N*, x-axis), and number of sampled time points (colour). Boxes: 10th, 50th, 90th percentiles. (B) Sankey diagram showing how filtering fits based on the sign of 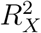 enriches for fits with positive accuracy against ground truth (*R*^2^ *>* 0). (C) Comparison of accuracy metrics (*ρ, r, R*^2^) against ground truth, stratified by the sign of the cross-validation score (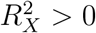vs.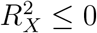). Boxes: 10th, 50th, 90th percentiles.

## 4 Application to Experimental Data

ALFA-K was also applied to real-world longitudinal single-cell DNA sequencing data to infer fitness landscapes and forecast evolution in experimental systems, as described in the main text. Here we detail the source and processing of this data.

### 4.1 Data Source and Experimental Systems

The experimental data analyzed in this study originates from the longitudinal single-cell copy number sequencing dataset published by Salehi et al. [23]. Their work characterized karyotype evolution in two experimental systems: immortalized human mammary epithelial cell lines (184-hTERT, including wild-type *TP53* ^WT^ and two independent *TP53* ^-/-^ lines) serially passaged for multiple generations, and four PDX models serially passaged in mice. Salehi and colleagues generated per-cell integer copy-number profiles across fixed genomic bins with the HMMCopy pipeline and applied the quality-control filters detailed in their publication [23].

Post-QC data were provided as a cell by genomic-bin integer matrix. A small number of samples were omitted because their reported genomic loci were inconsistent with the reference coordinates. Only bins from chromosomes 1–22 were retained. For each cell, the modal copy number across all bins assigned to the same chromosome was taken as that chromosome’s copy number, yielding a 22-element integer karyotype vector; sex chromosomes were ignored. Experimental populations in the Salehi et al. dataset were serially passaged and sampled at multiple time points. Within a population, we defined the chronologically succeeding sample as the daughter of the immediately preceding sample, and every chain of at least two successive samples was kept as a lineage. For every lineage, chromosome-level karyotypes were grouped by sampling passage, and the number of cells displaying each distinct karyotype was tallied. These karyotype-by-passage matrices, paired with an assumed passage interval of 15 days (PDX) or 5 days (184-hTERT), constituted the inputs for ALFA-K fitness-inference analyses.

**Figure S4:**
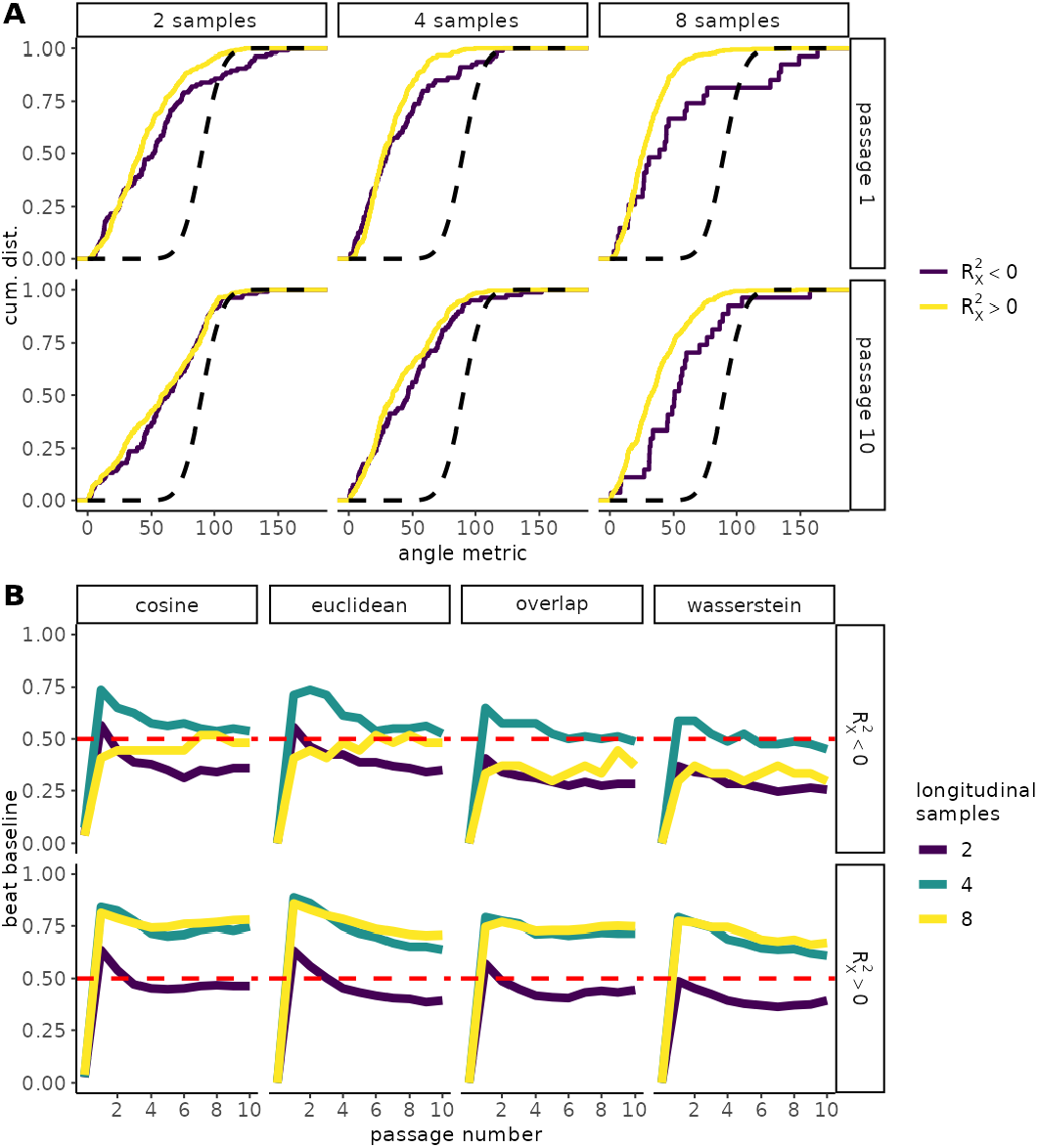
Evaluating the forecasting performance of ALFA-K fitness landscapes using ABM simulations. (**A**) Directional accuracy assessed by the angle metric *θ*. Empirical cumulative distribution functions (ECDFs) of *θ* are shown, faceted by number of training samples (columns) and forecast horizon (rows, in passages). Lines show the theoretical null distribution (black dashed) and results for ALFA-K fits stratified by cross-validation score (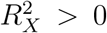vs.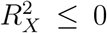, colored lines). Left-shifted ECDFs indicate better directional accuracy. (**B**) Performance relative to no-evolution baselines. Facets show the fraction of ALFA-K forecasts outperforming a given baseline metric (columns; see Section 3.1) stratified by cross-validation score (rows). X-axis is the forecast horizon (passages); colored lines indicate number of training samples. Higher fractions indicate better relative performance for ALFA-K.

### 4.2 Modeling the Influence of Karyotype Background on Fitness Effects

#### Data Processing and Trajectory Selection

Longitudinal copy–number data were expressed as a series of <u>transitions</u>, each defined by two consecutive population samples. Every ALFA-K fit corresponds to a trajectory—an ordered list of these transitions—along with its length (*n*_*trans*_). Within each bootstrap iteration trajectories were selected at random with probability proportional to (*n*_*trans*_) and retained only if no transitions had appeared in a trajectory already chosen. This was repeated until no more non-overlapping trajectories remained. This procedure maximises coverage of long evolutionary paths while ensuring that each transition is analysed exactly once. Trajectories containing a single transition and those with negative CV scores were excluded.

#### Modeling Fitness Effect Distributions (GLMMs)

Generalized Linear Mixed Models (GLMMs) were employed to analyze how context and treatment affect the properties of the Δ*f* distributions, specifically their magnitude and variance, while accounting for the nested structure of the data using the glmmTMB R package.

#### Model for Absolute Fitness Effect Magnitude (|Δ*f*|)

Let *R*_*ijk*_ be the scaled absolute fitness effect (|Δ*f*| + 10^−6^) for the *k*-th potential mutation relative to the *j*-th focal karyotype within the *i*-th trajectory fit. The model treats *R*_*ijk*_ as Gamma distributed with mean *µ*_*ijk*_ and dispersion *ϕ*:

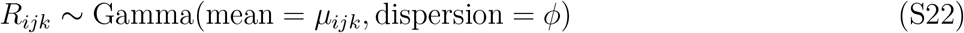

The relationship between the mean and the fixed and random effects is modeled via a log link:

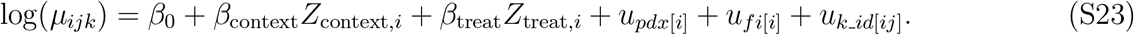

Where:

- *β*_0_ is the overall intercept.
- *β*_context_ and *β*_treat_ are the fixed effects coefficients for context (PDX vs. in vitro) and treatment (cisplatin vs. control, with cisplatin as the baseline level). *Z* denotes the corresponding indicator variables.
- 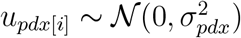is the random intercept for the PDX/cell line associated with trajectory *i*.
- 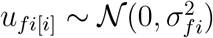 is the random intercept for trajectory *i*, nested within PDX line.
- 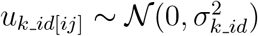 is the random intercept for the *j*-th focal karyotype, nested within trajectory *i*.

This model structure was fitted separately testing ‘context’ and ‘treat’ predictors (for ‘treat’, only PDX data was used).

#### Model for Variance of Fitness Effects (log(Var(Δ*f*)))

Let *V*_*ij*_ be the variance of the Δ*f* values across all potential mutations relative to the *j*-th focal karyotype within the *i*-th trajectory fit. The response variable is *U*_*ij*_ = log(*V*_*ij*_ + 10^−6^), scaled. This model treats *U*_*ij*_ as normally distributed:

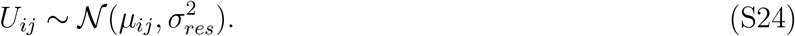

The mean *µ*_*ij*_ is modeled linearly:

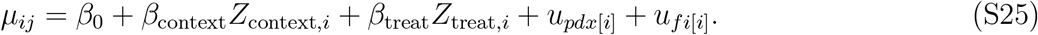

Where terms are defined similarly to the prior model (Eq. S23), but random effects 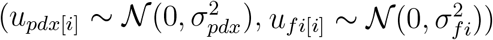 are only needed up to the trajectory level as each data point *U*_*ij*_ represents one karyotype.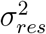 is the residual variance.

#### Estimating Fitness-Landscape Correlation Length (LMM)

Following the definition of correlation length in fitness landscapes formulated by [40], a long correlation length indicates a smooth landscape in which many single-chromosome changes are required to randomize fitness, while short correlation length characterizes a highly rugged landscape. To estimate this quantity we performed pairwise comparisons between the Δ*f* vectors (*v*_*p*_, *v*_*q*_) of different karyotypes (*p, q*). The Pearson correlation (*sim*_*pq*_ = cor(*v*_*p*_, *v*_*q*_)) and Manhattan distance (*dk*_*pq*_) were calculated. A Linear Mixed Model (LMM) was used to assess how similarity decays with distance using the <mosospace>lme4</mosospace> R package. The correlation *sim*_*pq*_ was transformed using the Fisher-z transformation (*G*_*pq*_ = atanh(*sim*_*pq*_)) to stabilize variance and approximate normality. The distance *dk*_*pq*_ was transformed using log(1 + *dk*_*pq*_) to potentially linearize the relationship and handle *dk* = 0. The model structure was:

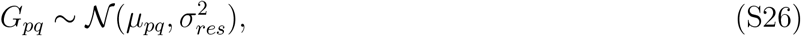

with:

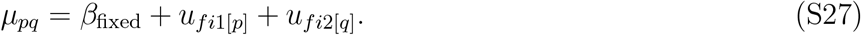

Where *β*_fixed_ represents the combined fixed effects part including main effects for log(1 +*dk*_*pq*_), pair type (same trajectory, parallel, different line), treatment pairing, and interaction terms between distance and pair type. Random intercepts 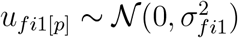 and 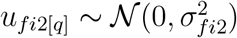 accounted for non-independence arising from karyotypes belonging to specific trajectories.

#### Trajectory Dynamics

The overlap in clonal composition between successive passages (sum of minimum frequencies) and the angular similarity between vectors of karyotypic change across passages were calculated to assess the rate and directionality of evolution.

#### Modeling of Whole-Genome Doubling (WGD) Effects

Karyotypes were classified as WGD^+^ if their modal chromosome copy number was ≥ 3, and WGD^−^ otherwise. Aneuploidy level for a karyotype **k** was quantified as the number of altered chromosomes, *d*(**k**) = ∑ _*i*_ I(*k*_*i*_*≠* mode(**k**)), where *k*_*i*_ is the copy number of chromosome *i*.

#### Aneuploidy Divergence

The dynamics of aneuploidy accumulation over passages were modeled using non-linear least squares (via R’s `nls` function), fitting the average number of altered chromosomes per cell (*n*_*a*_) at passage *t* to an exponential saturation curve, *n*_*a*_ = *A*(1 − *e*^−*κt*^). Both the asymptote parameter (*A*) and the rate constant parameter (*κ*) were modeled using additive fixed effects for WGD status (*s* ∈ {WGD^−^, WGD^+^}) and experimental trajectory (*j* ∈ {TrajA, TrajB}), assuming no interaction: *A*_*sj*_ = *β*_*A*0_ + *β*_*A,W GD*_ *×* 𝕀 (*s* = WGD^+^) + *β*_*A,T raj*_ *×* 𝕀 (*j* = TrajB) and *κ*_*sj*_ = *β*_*κ*0_ + *β*_*κ,W GD*_ *×* 𝕀 (*s* = WGD^+^) + *β*_*κ,T raj*_ *×* 𝕀 (*j* = TrajB), where I(*·*) is the indicator function.

#### Fitness Effect Distributions

Distributions of single-chromosome gain/loss fitness effects (Δ*f*) for WGD^+^ vs WGD^−^ states were compared using a permutation Kolmogorov-Smirnov test. Mean effects were calculated per WGD status within each landscape fit before comparison. Significance was assessed against 10^4^ permutations of WGD labels among fits (*P <* 0.0001).

